# Bayesian methods for optimizing deep brain stimulation to enhance cognitive control

**DOI:** 10.1101/2022.12.14.520473

**Authors:** Sumedh S Nagrale, Ali Yousefi, Theoden I Netoff, Alik S Widge

## Abstract

**Objective:** Deep brain stimulation (DBS) of the ventral internal capsule/striatum (VCVS) is a potentially effective treatment for several mental health disorders when conventional therapeutics fail. Its effectiveness, however, depends on correct programming to engage VCVS sub-circuits. VCVS programming is currently an iterative, time-consuming process, with weeks between setting changes and reliance on noisy, subjective self-reports. An objective measure of circuit engagement might allow individual settings to be tested in seconds to minutes, reducing the time to response and increasing patient and clinician confidence in the chosen settings. Here, we present an approach to measuring and optimizing that circuit engagement.

**Approach:** We leverage prior results showing that effective VCVS DBS engages circuits of cognitive control, that this engagement depends primarily on which contact(s) are activated, and that circuit engagement can be tracked through a state space modeling framework. We combine this framework with an adaptive optimizer to perform a principled exploration of electrode contacts and identify the contacts that maximally improve cognitive control.

**Main results:** Using behavioral simulations directly derived from patient data, we show that an Upper Confidence Bound (UCB1) algorithm outperforms other optimizers (roughly 80% probability of convergence to a global optimum).

**Significance:** We show that the optimization can converge even with lag between stimulation and effect, and that a complete optimization can be done in a clinically feasible timespan (a few hours). Further, the approach requires no specialized recording or imaging hardware, and thus could be a scalable path to expand the use of DBS in psychiatric and other non-motor applications.

## Introduction

Deep brain stimulation (DBS) is a well established, clinically successful surgical treatment for medication-refractory movement disorders[1]. It is being explored as a treatment for mental disorders, including obsessive compulsive disorder (OCD), major depression (MDD), addiction, and schizophrenia[2–5], in cases where conventional therapies such as medication and psychotherapy have failed. The ventral internal capsule/ventral striatum (VCVS) is a particularly promising DBS target. It is approved by the US Food and Drug Administration for treatment of OCD and is the only psychiatric DBS target to meet its endpoint in a randomized controlled trial for MDD [6]. At the same time, VCVS DBS consistently shows a 60-70% response rate in large open label studies [7–9], meaning that roughly 1/3 of patients undergo this invasive intervention for little benefit, and another 1/3 are left with substantial, life-impairing residual symptoms.

These difficulties arise in part because the success of VCVS DBS depends upon proper programming. The only successful VCVS DBS trial in MDD used an extended programming period, with a year of open-label treatment and adjustment before blinded testing[6]. DBS programming is challenging regardless of the target or disorder. The large number of free parameters (which contacts are active at which polarity, waveform amplitude, pulse width, and stimulation frequency) creates a combinatorially explosive space. Worse, brain responses to stimulation are often non-linear[10–12]. In tremor-related disorders (the primary clinical application for DBS), this challenge is partly overcome by the speed of response. Effective settings cause tremor changes within seconds, providing immediate feedback to help a clinician identify reasonable parameters. In mental disorders, while there are immediate subjective effects of some setting changes[13,14], these do not consistently predict long-term outcomes[2,15,16]. Rather, programming is usually based on patient self-report of symptoms over days to weeks[8,15,16]. Self report has a poor signal-to-noise ratio, as patients’ symptoms can be heavily influenced by external events (e.g., mood and compulsive symptoms being dramatically worsened by the recent pandemic[17]) and the subject stimulation response can depend on those externally-influenced states[18]. The long latency between setting changes makes it much more difficult to fully explore the DBS parameter space. Thus, much of the apparent non-response to VCVS DBS, across disorders, may represent a target engagement problem – some patients may never receive stimulation that adequately affects the brain circuits mediating clinical response[2,19,20].

Recent work, at VCVS and other targets[21–23], argues that target engagement could be resolved through imaging. In this view, the correct contact and other settings can be identified based on electric field modeling, with the goal being to electrically engage specific white matter bundles. More recent work, however, finds that imaging-based target engagement does not fully predict or determine clinical response[24,25]. This suggests that, beyond anatomy, the programmed parameters need to cause a specific physiologic response in the target and its connected network to achieve clinical response. In theory, closed loop sensing, e.g. of the local field potential at the DBS target or a connected structure, might allow monitoring of that physiologic response[26–29]. This approach has been successful in movement disorders[30–32]. The challenge is that, again owing to the poor signal-to-noise ratio of self report, it has been difficult to identify reliable physiologic markers for DBS tuning[27,28,33]. It may be possible to derive such markers by extensive data collection and patient-specific model fitting[18,26,34], but the analytic pipelines required for such modeling may not scale well to clinical practice.

A more scalable approach might be to measure physiologic engagement indirectly, through effects on behavior. Changes in clinically relevant circuits should be reflected in relatively rapid changes in brain functions linked to those circuits[35]. Just as DBS for tremor can be rapidly titrated by monitoring tremor in-office, it should be possible to continuously monitor an objective behavior (e.g., performance on a psychophysical task) and titrate stimulation to optimize that behavior. We recently demonstrated a prototype of that approach. In two human studies, we showed that DBS-frequency stimulation of VCVS enhances cognitive control[36,37]. Cognitive control is the ability to withhold a prepotent response in favor of a more adaptive, long-term-goal-oriented response. It is an attractive target for measurement and enhancement with DBS, since cognitive control deficits are common across a range of mental disorders[38–40]. A method for engaging cognitive control circuits could thus make VCVS DBS applicable for a wide range of clinical problems. We showed that cognitive control could be measured through DBS-induced changes in reaction time on a standard cognitive conflict task, the Multi Source Interference Task (MSIT)[36,37]. Participants who experienced stimulation-induced control enhancement also reported improved depressive and anxious symptoms. Most importantly, these improvements were sensitive to stimulation location within the VCVS. We saw generally larger effects with right-sided stimulation in the more dorsal capsule, but the best location varied between patients. Further, the behavioral effects of stimulation began and ended within a few seconds of stimulation changes[37]. It should thus be possible to use these rapid-onset effects to identify when the optimal contact is being stimulated. The challenge is that because the reaction time effects are subtle (5-10% of the overall scale), they cannot be immediately detected by a human operator, nor can the effects of different contacts be quickly distinguished. These subtle changes can be detected by a monitoring algorithm, if it uses a filtering/smoothing approach to ignore stochastic reaction time variability[37]. The missing ingredient is an approach for using that monitoring to quickly but reliably identify the optimal stimulation site and other parameters for engaging cognitive control circuitry.

Here, we present such an approach, based on Bayesian optimization. Bayesian optimizers maintain a prior estimate of the relative effects of different parameters (e.g., of different contacts, or of changes in amplitude within a contact). They then select a next contact to test, based on an information gain function that varies with the specific optimizer (see examples in Methods). Based on the observed behavioral response, the prior estimate is updated (posterior distribution calculation), then the next test value is selected. The process repeats until a convergence criterion is met. Bayesian methods have been suggested as an efficient way for tuning DBS parameters in Parkinson disease [41], and we recently demonstrated their value in tuning neurostimulation for pain [42]. Using a behavioral simulator derived from our prior empirical data, we show that a specific optimizer (Upper Confidence Bound) and rate of change in stimulation parameters can reliably converge to a global optimum, the vast majority of the time, in a clinically feasible amount of testing.

## Methods

### Overall design

Testing a wide range of optimization strategies directly in patients undergoing VCVS DBS is impractical. It would require thousands of task trials across hundreds of runs, which is unlikely to be tolerated by most patients. Thus, we developed a simulation testbed for our cognitive control paradigm. The simulation is meant to model a patient continuously performing the MSIT (or a similar cognitive control assay), while stimulation is adjusted to improve task performance (to minimize reaction times without inducing errors). The overall system (Figure 1A) includes a data generator, a sensor, and an optimizer. The generator models a patient performing the MSIT and emitting behavioral data, in the form of reaction time (RT) values that are simultaneously influenced by stimulation, task factors, and noise processes. The sensor models the real-time estimation we would then need to do with a patient, analogous to the processing in [37]. It attempts to filter out the noise and task processes to identify stimulation-driven RT changes, without explicit information about the current stimulation settings. This smoothed derivative of RT (sometimes referred to as a “cognitive state”) is then used by the optimization algorithm to update its distribution estimate. Based on the update, the optimizer then modifies the simulated stimulation applied to the generator. This process continues until the algorithm converges.

**Figure 1:**
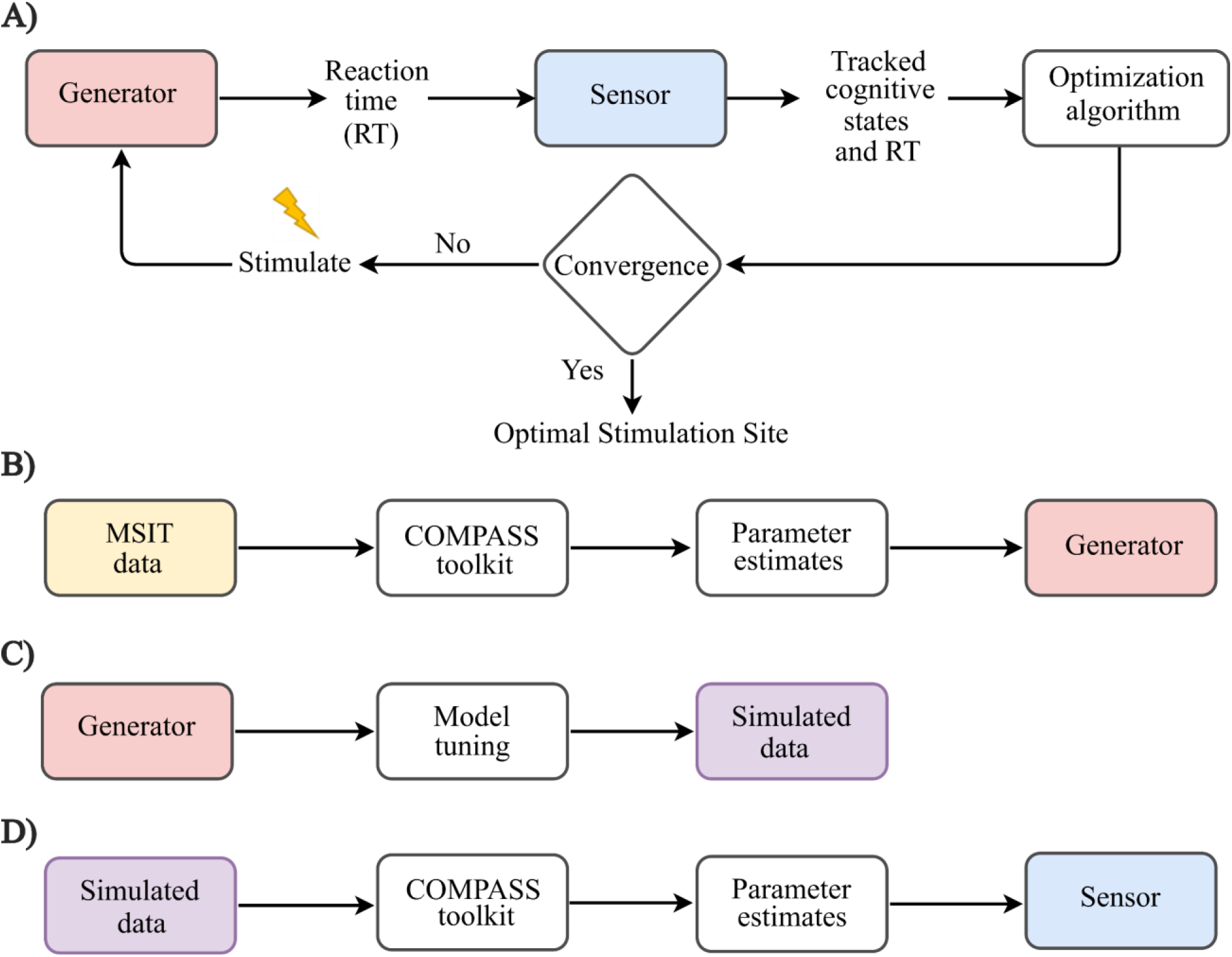
System design. A) The generator model imitates a DBS recipient. A sensor model infers the generator’s internal/unobservable cognitive state based on the observable reaction time (RT). The optimization algorithm then takes the cognitive state estimate as input and selects a new stimulation regime to be applied to the generator. Over time, the algorithms converge and the optimal or near optimal location is determined. B) The internal parameters of the generator are inferred by model fitting to data from actual participants who performed MSIT and received stimulation. C) Data is simulated using the generator model and a gamma distribution to approximate human RTs. New simulated patients can be created by slightly modifying the internal generator parameters or initial state (“model tuning”). D) The sensor model is derived by fitting a new model to a block of simulated RT data from the generator. Once derived, it continuously estimates the generator’s internal state trial-by-trial through a Kalman-like filtering process[43].

In the current work, we focused on the problem described in [37], optimizing the stimulation location (choosing among available DBS contacts or combinations of contacts), assuming that other parameters such as amplitude, pulse width, and frequency are already reasonable. We had empirical data to constrain this problem (see below), making it a logical starting point.

### Empirical data

The dataset includes 6 participants with long-standing pharmaco-resistant epilepsy. They were reported as participants 8-13 in [37]. We relabel them here as S1-S6. Participants voluntarily enrolled with fully informed consent (obtained by a member of the study staff who was not the participant’s primary clinician) in accordance with guidelines and procedures approved by the local institutional review boards at Partners Healthcare (Massachusetts General Hospital), with secondary review from the US Army Human Research Protections Office.

Participants performed the MSIT with simultaneous behavior and local field potential (LFP) recordings. Stimuli for the MSIT were presented on a computer screen with either Presentation software (Neurobehavioral Systems) or Psychophysics Toolbox. Each participant performed 1 to 3 sessions, and each session consisted of multiple blocks of 32 or 64 trials, (0.9 to 57.6 min) with brief rest periods (median break time of 5.25 min) in between the blocks. They were instructed to be as fast and as accurate as possible. Median success rates of 100% ± 2.47% and 97.1 ± 5.52% were reported during congruent and incongruent conditions, respectively[37]. The task contained a roughly equal number of congruent and incongruent trials. Stimuli were presented for 1.75 s, with an inter-trial interval randomly jittered within 2–4 s. Simultaneously, LFPs were recorded using implanted depth electrodes; these data were not considered in the current study. The electrode locations were determined by a team of caregivers solely on clinical grounds and were not in any way modified for the research.

Only one site was stimulated during each MSIT block. Stimulation was a 600ms long train of symmetric biphasic (charge balanced) 2–4mA, with 90μs square pulses at a frequency of 130 Hz, delivered using a Cerestim 96 (Blackrock Instruments). Stimulation was delivered through a neighboring pair of contacts on a single depth electrode (bipolar) with parameters set manually by the experimenter. Depth electrodes (Ad-Tech Medical or PMT) had a diameter of 0.8-1.0 mm and had 8-16 contacts (platinum/iridium), each 1-2.4mm long. The stimulations were triggered by a separate PC that was either delivering or monitoring the task or behavioral stimuli. All stimulations were delivered at the image onset to influence a decision-making process that begins with that onset. Participants S1-S6 performed 343, 378, 320, 383, 383, and 440 trials respectively. This included both stimulation and non-stimulation trials. All stimulations were given to varying sites within the internal capsule as described in [37].

### Data model

The MSIT is described extensively in [36,37]. In brief, it is a standard cognitive control/conflict paradigm where roughly 50% of trials contain distractors that induce pre-potent (incorrect) responses. These are referred to as “incongruent”, “interference”, or “high conflict” trials. Successful task performance requires suppression of those responses, both through reactive processes (e.g., stopping an incorrect action once started) and proactive processes (e.g., engaging attentional systems to detect impending errors). In this context, task RTs measure a participant’s ability to apply cognitive control – faster RTs without errors imply more efficient control systems.

Task reaction times (RTs) are positive random variables with right-skewed distributions[44,45]. In particular, RTs are often well described by Gamma distributions, with parameters that vary depending on the trial type (e.g., the high vs. low conflict trials in MSIT). Thus, we model the reaction time of a participant performing MSIT as described in [45],

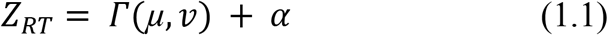

Where Z_RT_ is the observed reaction time, α is the offset term representing the lower bound of the reaction time, 1/v is the dispersion observed in the reaction time, and μ is the mean RT.

That mean RT varies both systematically and stochastically. Systematically, it is higher on high conflict trials. Stochastically, it has both random variation (captured in *v*) and slow change over time. The latter may be due to learning processes (participants get slightly better at the task over the first 100-200 trials) or, more commonly, due to the effect of brain stimulation that alters the participant’ s “baseline” RT. As in [37], we express this in terms of two unobservable “cognitive states”, *x_base_* and *x_conflict_*, connected to the RT by a log link function[45,46],

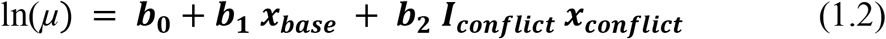

*x_base_* is the baseline cognitive state, which represents the mean RT when no conflict is present. *x_conflict_* is the mean amount of increase in RT during high conflict. ***I_conflict_*** is an indicator variable representing presence or absence of conflict in a given trial of MSIT. b_0_, b_1_, b_2_ are then weights (effectively, regression coefficients) that relate cognitive states to the distribution mean μ. b0 provides a bound on RT, whereas b_1_, b_2_ are scaling factors. We have demonstrated this model as well suited to MSIT datasets in multiple prior papers[37,43,46].

The temporal evolution of cognitive states at trial *k* is defined by a first order autoregressive model, AR(1), as in [45,46].

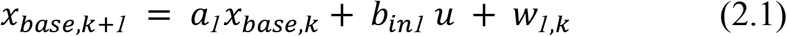

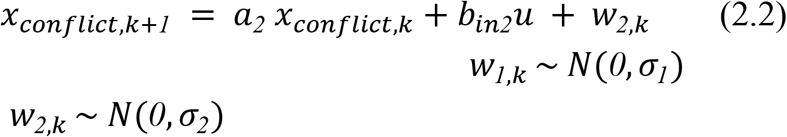

The next states *x_base,k+1_*, *x_conflict,k+1_* are dependent on the current states *x_base,k_*, *x_conflict,k_*, the input u (which represents the presence of applied DBS), and the Gaussian state noise processes *w_1,k_*, *w_2,k_* with zero mean and variance σ_1_, σ_2_ respectively.

Here, we simulated a case analogous to our empirical experiments in [37], where we tested different contacts along a cylindrical lead while holding other stimulation parameters (amplitude, pulse width) constant. In this simulation, the input vector u has length equal to the number of available contacts, all of which are assumed to be stimulated in monopolar mode (as in a clinical monopolar survey). The elements of u are 1 when a given contact is active, and 0 when it is not. The weighting vectors *b_in1_*, *b_in2_*then describe how stimulation alters the state variables. Note that in this formulation, effective DBS changes these underlying generating states, but does not act directly on the mean RT *μ*. Thus, when a new stimulation regime is applied, the states *x_base_* and *x_conflict_* will not immediately change but will drift to a new steady state. This is consistent with our observations in [37]. The rate of drift will depend on *a* and *b_in_*. As further described below, this influences the design of the optimization approach.

### Generator model

The state space model of (1) and (2) is applied to generate new samples of behavior (MSIT RTs) that model those we would expect to observe from a patient undergoing DBS. The evolution of behavior is represented by stepping (2.1) and (2.2) forward in time. The modeling requires identifying reasonable values for the parameters in these equations.

We perform parameter and state estimation using the COMPASS toolkit[43], which uses a combination of an Expectation Maximization (EM) algorithm and a Kalman-type filtering/smoothing approach to simultaneously infer model parameters and cognitive states given the observed behavior in a block of trials.

The parameter estimation problem for MSIT as described in (1) and (2) would be

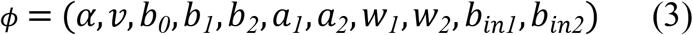

This forms a complex estimation problem with a large number of unknown parameters This increases the possibility of converging to a local maximum, resulting in sub optimal estimates. To prevent this, we reduce the parameter estimation search space. First, we temporarily ignore the input weightings *b_in1_* and *b_in2_*, by setting them to 0. As described below in section “Optimizer test with simulated stimulation response surfaces”, we will assign new values to these parameters when performing the simulation, making it irrelevant to estimate them from patients’ data. Note that we can still use data from patients receiving stimulation (as in [37]) to estimate the model – any drift in *x_base_* and *x_conflict_* caused by stimulation is captured by the dispersion 1/v and and state Gaussian noise *W*. For convenience, here the generator models were estimated only from non-stimulation blocks within the empirical dataset.

As a further simplification, we apply no scaling to the cognitive states, which is done by setting scaling parameters *b_1_* and *b_2_* to 1. Finally, we recognize that the state noise process *w* is the most challenging free parameter to estimate, as a wide range of values can be consistent with observed data. We therefore assign values to *σ_1_* and *σ_2_* that we identify as working well across patients; these are described further in “Grid search for model noise processes” below. This simplifies the parameter search space to

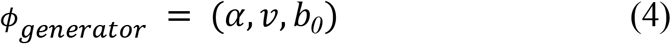

The problem (4) is then estimated using the COMPASS EM algorithm. The convergence of the EM algorithm is determined by assessing the maximum likelihood (ML) of the data at each iteration, given the current parameters *ϕ_generator_*. The convergence criterion for EM algorithm is a plateau in the ML change given by (4.a)

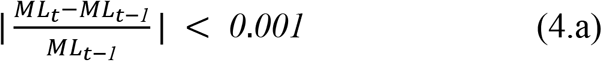

To demonstrate the models’ ability to simulate new participants that are very similar to the empirical data, 1000 new models (“generator pool”) were estimated based on each participant’s data (S1-S6). For each model fit, the initial conditions *x_0_* in (2) were selected randomly from a normal distribution N(0,1), autoregressive parameters *a* were selected close to 1 [*a_1_*=0.9999, *a_2_*=0.9999] and for *W* values for *σ_1_* and *σ_2_* were randomly selected from the scale range [7,16] and [13,27] respectively identified in “Model noise process determination via grid search”.

### Generator validation

To validate the generator model, we computed the Kolmogorov-Smirnov (KS) distance between the empirical distribution of those simulated RTs and the empirical probability distribution function (pdf) of a gamma distribution fit to the simulated RTs (MATLAB fitdist function). A KS distance/test at p>0.05 would suggest that the generator is producing RT-like, gamma-distributed data. We performed two versions of the simulation, one with model tuning and other without model tuning. Model tuning was performed to demonstrate that, if necessary, the generator model could closely replicate the actual distribution obtained from any given empirical participant. Model tuning was a trial and error method of modifying model parameters A, v, w and *α* by hand. We simulated both the tuned model and untuned model each for 1000 trials as described above. These simulations randomly generated interference trials by setting *I_conflict_* to 0 or 1 on each trial. No stimulation effects were included. This was repeated 1000 times for each participant, followed by the KS test as just noted. We report the fraction of simulations that had p>0.05.

### Sensor model

In actual patients, we do not have direct access to their underlying cognitive states but infer these by fitting the model of (1-2) to the observed RTs. In our simulations, however, the ground truth is known, and we can confirm that our sensor model accurately recovers it. The sensor model fitting problem, analogous to the generator problem, is given by

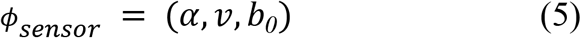

The difference between the sensor and the generator model is the data used for modeling and evaluation. For the generator model, we fit it and tested its deviance against the empirical data from S1-S6. For the sensor model, we used data simulated from a pool of randomly initialized generators. We used the same COMPASS EM algorithm, with convergence rule (4.a), and *σ_1_*, *σ_2_* drawn from the scale range [7,12] and [13,27]; the origin of this choice is described below in “Grid search for model noise processes”. Further, we added 0.0625 (equal to scale parameter 8) to both *σ_1_* and *σ_2_*. This slight increase in noise covariance is helpful in later modeling steps, when we attempt to capture the effect of stimulation. Because the sensor model does not explicitly represent stimulation, a slightly higher state noise leads to more rapid adaptation when stimulation is applied and the state variable is forced away from its default AR(1) behavior.

For each sensor model, the initial state *x_0_* in (2) was randomly selected from a normal distribution N(0,1). To further accommodate a wide range of generators, we set the lower bound *α* of reaction time to 0.1. We simulated 1000 fits of a sensor model to a generator model. The generator for each simulation was chosen from the “generator pool” described above under “Generator model”. We then evaluated sensor tracking ability on this randomly selected generator model using normalized root mean squared error (NRMSE), as described in “Sensor tracking validation” below.

### Grid search for model noise processes

For the problem described in (3), a large number of unknown parameters increases the probability of incorrect convergence of the EM algorithm. In initial pilots, we noticed that this was particularly problematic when attempting to estimate the covariance of the state noise process *W*. (Here, *W* is a diagonal matrix containing parameters *σ_1_, σ_2_*). This estimation often converged to different values, with diagonal elements between [*10^−1^*, *10^−7^*]. All other parameters in *ϕ* would be altered in turn. This wide range of *W* values is not reasonable on face value, suggesting a need for a constraint. We theorized that there might be acceptable values of *W* that may work well across all patients, simplifying the problem of (3) to (4) and (5).

We tested this theory by a two-dimensional grid search, using a logarithmically spaced grid. At each grid point, we set the diagonal elements of *W* to the specified values, estimated *ϕ_generator_* using COMPASS, then calculated the deviance between observed RTs and those predicted by the model (μ from equation 1.2). The logarithmic grid is given by

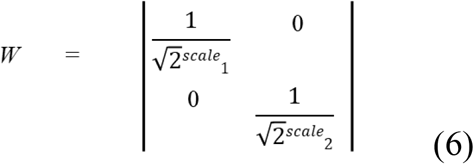

Where *scale_1_ ∊* [1,40] and *scale_2_ \** [1,40]. The range was selected empirically based on values observed in previous studies [37]. Numerically, *σ_1_*, *σ_2_* takes values from [9.5367e-07, 0.707] for logarithmic search. For each point in the grid search, the problem (4) was estimated by fitting to data including both non-stimulation and stimulation trials from the empirical dataset using the COMPASS EM algorithm. We then evaluated the deviance between observed RTs and the one predicted by the model. This fit was performed separately for each dataset for participants S1-S6. It was repeated 5 times for each participant, using a random *x_0_* selected from N(0,1), plus an additional run with initial condition *x_0_* set to 0. For this model inference we set *A_k_* close to 1 [*a_1_*=0.9999, *a_2_*=0.9999], *W* was set based on the current grid point, b was 0 and the overall distribution was assumed to be gamma.

### Sensor tracking validation

In previous reports, we validated sensor models by showing that the residuals of predicted RTs were consistent with a white noise process[37,46]. Here, because we had access to the ground truth generator internal states, we directly measured whether the sensor model accurately tracked those states. We calculated the normalized root mean square error (NRMSE) between the *x_base_* and *x_conflict_* values in the generator and those inferred by the sensor. NRMSE is the root mean square error (RMSE) between the generator and sensor states after both variables have been re-scaled multiplicatively to the [-1,1] interval. Because our optimization is concerned primarily with changes in a state variable relative to its own baseline, and not with the specific numeric value of that variable, NRMSE is a more appropriate metric. It tracks whether the sensor’s estimate changes in the same direction and scale as the generator ground truth.

NRMSE was calculated between the sensor model (predicted RT and cognitive states) and the corresponding ground truth from the generator. We selected generators randomly from the generator pool (1000 models from 6 participants as noted above). To ensure that we demonstrated that the sensors tracked well over all assumed noise process values, we enforced a further constraint that *σ_1_* in the generator be drawn uniformly from the selected scale range of [7, 12]. The generator-sensor models were simulated for 1000 trials each, for 1000 randomly selected generators and sensors. During these trials, interference variable *I_conflict_* was set randomly on each trial, and the value of this indicator was visible to both sensor and generator.

To illustrate the sensor’s timescale for settling to a steady state when stimulation is changed, we also simulated the two-site, high contrast stimulation *b_ss_* as described in “Optimizer test with simulated stimulation response surfaces”. This simulation also used 1000 trials, but stimulation was changed every 25 trials between the two elements of *b_ss_*.

### Bayesian Optimization to identify optimal stimulation site

Identifying the optimal stimulation contact on a cylindrical DBS lead, assuming that other parameters are held constant during this survey, is a multi arm bandit (MAB) problem. MABs consist of competing discrete options (here, k independent stimulation sites from available stimulation sites in u), of which one of the options produces the maximum effect (here, reduction in the RT). Common algorithms for solving MABs include a greedy algorithm [47,48], *∊*-greedy algorithm [47,48], Upper Confidence Bound (UCB1)[47], Bayesian upper confidence bound (Bayes-UCB) [49]and Thompson sampling (TS)[48].

In general, the goal of these algorithms is to minimize the approximate cost function *Q*(*u*) by performing a trade off between exploring (selecting arms with uncertain payoff) and exploiting (selecting the arm currently estimated to have the best payoff). Algorithms differ in the cost functions and the steps they take to perform the trade off between exploration-exploitation, as described below.

#### Greedy and e-greedy algorithm

The pure greedy algorithm solely exploits and is a degenerate case of *∊*-greedy with *∊*=0. The ε-greedy algorithm is an improvement to the greedy algorithm which primarily exploits, but also explores a new stimulation site *u_k_* randomly with a small probability *∊* (here selected as 0.1). The cost function *Q*(*u*) is adapted from [47]

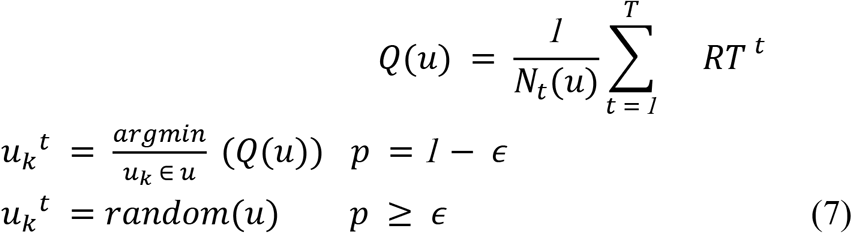

Where *u* is the input vector to (2), with 1 indicating the selected site *u_k_*, *N_t_*(*u*) is the number of times a site is selected, T is the overall number of trials done so far, *p* is a random probability drawn uniformly from [0,1], and *u_k_^t^* is the stimulation site k selected at time t based on cost function.

#### Upper Confidence Bound (UCB1)

Upper Confidence Bound (UCB1) explores and exploits based on the Hoeffding inequality [47] that accounts for the number of times a particular stimulation site *u_k_* has been selected. It maintains an estimate of each site’s worst-case performance, with a confidence interval around that estimate that shrinks as the site is selected more. Over time, as all sites have been repeatedly selected, UCB1 becomes less exploratory and more exploitative. The cost function [47] for UCB1 is

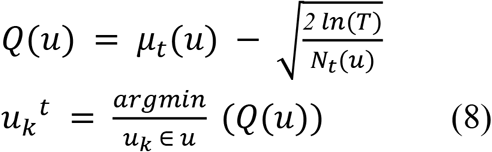

Where *μ_t_*(*u*) is the vector of the mean RT of stimulation sites in u.

#### Bayesian inference and Conjugate prior

Bayesian algorithms differ from UCB1 in that UCB1 directly maintains an estimate of the distribution of observed RTs, whereas Bayesian algorithms maintain an estimate of the parameters defining that distribution.

The exploration-exploitation of stimulation sites is based on these belief dynamics about the quality of each available site. This belief is expressed in terms of prior *П^0^* and posterior distributions *П^t^*. The *П^0^* is the probability distribution before the RT is observed at t, whereas the *П^t^* is the probability distribution after observation at t. On each trial t, a random hypothetical RT *h^t^* is drawn for each stimulation site *u_k_* in u, using the prior estimate of the distribution parameters *θ_k_*, and the site with the best draw is selected by (9)

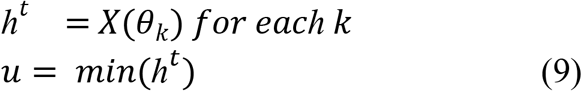

The Bayesian algorithms assume the RT observed on each trial to be an independent identically distributed (IID) draw. With each observation, they perform a posterior update of the parameter belief to obtain a posterior belief *П^t^*, following Bayes’ rule. This *П^t^* then is the prior for the subsequent trial at *t* + 1.

Prior and posterior distributions are selected from conjugate pairs. For conjugate distributions the prior (conjugate prior) and the posterior distribution fall in the same probability distribution family [48]. Here, the posterior update is performed in conjunction with a rule described in (10), which transforms the observed RT into a reward/payoff metric for the action of selecting stimulation site *u_k_*.

For any arbitrary distribution parameters *θ_1_*, *θ_2_* ∈ *θ_n_*,

if *RT_k_^t^* ≥ *RT_val_*: update *θ_1_*;

else: update *θ_2_*

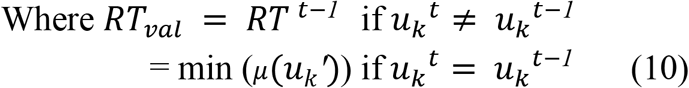

Where 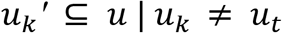. (10) states that if the current and previously selected stimulation site are different, then the threshold *RT_val_* is the previous *RT^t−1^* Otherwise, *RT_val_* is the minimum RT among the mean RTs of the rest of the stimulation sites. The parameters *θ_1_* and *θ_2_* depend on the specific choice of conjugate prior. The relevant distributions and their update rules are expressed in Table 1.

**Table 1:** Conjugate prior and posterior distributions and update rules for Bayesian multi-arm bandit algorithms.

| Algorithms | Likelihood | Conjugate Prior $\Pi^0$ | Posterior hyperparameters update $\Pi^t$ | |
| --- | --- | --- | --- | --- |
| | | | RT $\geq X$ | RT $< X$ |
| Bayes-UCB | Normal<br>RT$\sim N(\mu, \sigma^2)$ | Gamma<br>($\alpha, \beta$) | $\alpha = \alpha + \frac{n}{2}$ | $\beta = \beta + \frac{\sum_{t=1}^T (RT_t - \mu)^2}{2}$ |
| Bernoulli-TS | Bernoulli<br>$r_t \sim \text{Ber}(p)$ | Beta($\alpha, \beta$) | $\alpha = \alpha + r_t$<br>$r_t \in [0, 1]$ | $\beta = \beta + 1 - r_t$<br>$r_t \in [0, 1]$ |
| TS-Poisson | Poisson<br>$r_t \sim \text{Pois}(\lambda)$ | Gamma<br>($\alpha, \beta$) | $\alpha = \alpha + RT^t$ | $\beta = \beta + 1 - r_t$<br>$r_t \in [0, 1]$ |
| TS-Normal | Normal<br>RT$\sim N(\mu, \frac{1}{\tau})$ | Gamma<br>($\alpha, \beta$) | $\alpha = \alpha + \frac{n}{2}$ | $\beta = \beta + \frac{\sum_{t=1}^T (RT_t - \mu)^2}{2}$ |
| C-TS | Normal<br>RT$\sim N(\mu, \sigma^2)$ | Normal<br>($\mu_z, \frac{1}{1+k_z}$) | $\mu_z = \frac{\mu_z k_z + RT^t}{k_z + 1}$ | $k_z = k_z + 1$ |

#### Bayesian UCB

Bayes-UCB assumes that the process generating the observed RT is normally distributed, where the parameters mean *μ* and variance *σ* for that distribution are unknown. A gamma distribution *Γ*(*α,β*) is an appropriate choice of conjugate prior (*П^0^*) for such a normally distributed process[49]. Prior to each trial *t*, Bayes-UCB estimates *μ* and *σ* using the prior shape (*α*) and rate (*β*) hyperparameters (11). It then calculates a cost function/hypothetical RT *h^t^* to select the best stimulation according to the current belief. Once the actual RT is observed, as per Bayes theorem the prior belief is updated using posterior update 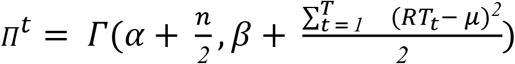 in combination with (10).

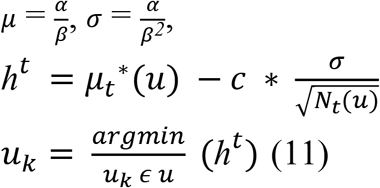

Where *c* is a confidence level value (here selected as 1.96 corresponding to 95% confidence level). The parameter *c* also adds to the exploration-exploitation. The margin of error 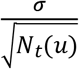 decreases as the sample size increases and algorithm is more certain about the mean *μ_t_*(*u*).

#### Thompson Sampling

Thompson Sampling (TS) is a further class of Bayesian algorithms that can achieve high efficiency sampling of complex MAB problems[48]. We tested multiple variants of TS that differed in their assumptions about the payoff’s underlying generative distribution. These included Bernoulli, Poisson, and normal distributions.

The TS-Bernoulli algorithm assumes the payoff/reward *r_t_* (success/failure) for a selected stimulation site *k* is generated from a Bernoulli(*p_k_*) distribution. The payoff *r_t_* is the transformed reward from RT based on whether a selected stimulation site reduces RT or not. If it reduces, *r_t_* is 1 else 0. The unknown parameter *p_k_* describes the probability with which the selected stimulation site *k* reduces RT i.e. the distribution for *r_t_*. The choice of conjugate prior for a Bernoulli distributed process is *П^0^* = *Beta*(*α*, *β*), where *α* and *β* are the shape parameters. Prior to each trial TS-Bernoulli calculates hypothetical RT *h^t^* (12) from the beta distribution and once the observation is made, a stimulation site *k* is selected using (11). The posterior update *П^t^* = *Beta*(*α* + *r_t_*, *β* + 1 − *r_t_*) in combination with (10) is performed as shown in Table 1[48].

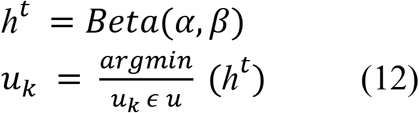

The TS-Poisson algorithm assumes the payoff/reward *r_t_* to follow a Poisson(*λ_k_*) distribution. The rate parameter *λ_k_* for each stimulation site *u_k_* captures the number of times *u_k_* is likely to reduce RT over the progression of the experiment. The choice of conjugate prior for a Poisson distributed process is *П^0^* = *Γ*(*α, β*) where *α* and *β* are the shape and rate parameters respectively. Before each trial t, TS-Poisson samples a hypothetical RT *h^t^* using the prior hyperparameters (13), then selects stimulation site k according to (11). Once RT is observed the hyperparameters are updated using posterior update *П^t^* = *Γ*(*α* + *RT^t^*, *β* + 1 − *r_t_*) in combination with (10) as shown in Table 1 [50].

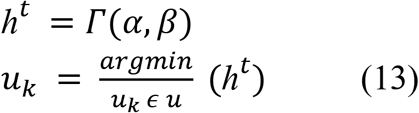

The TS-Normal algorithm (TS-Normal) assumes RT to follow a normal N(*μ, 1/τ*) distribution, with a known mean (*μ*) and unknown variance (1/*τ*), where *τ* is the precision. For a normal distribution with known mean, the convenient choice for conjugate prior is a Gamma distribution, *П^0^* = *Γ*(*α,β*) where *α* and *β* are the shape and rate parameters respectively. TS-Normal prior calculates hypothetical RT *h^t^*(14) directly using the prior hyperparameters and once the RT is observed, the posterior update is performed using 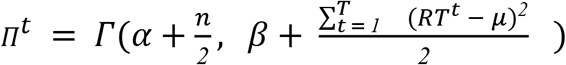 and (10) as shown in Table 1 with conjugate prior the same as Bayes-UCB[49]. The mean (*μ*) is updated with each iteration along with the posterior update.

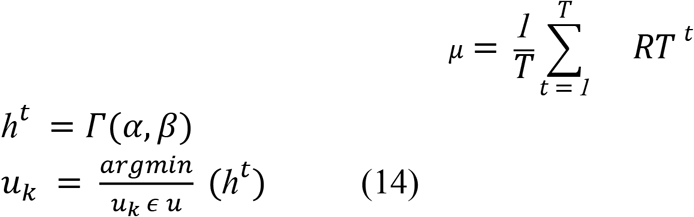

We adapted Causal Thompson sampling (C-TS)[51], for the stimulation site MAB problem. It assumes that the RT is produced from a Normal(*μ, σ*) distribution. Where the generator-sensor-optimizer is considered to be a causal model, and sequential intervention by selection stimulation site is considered to cause a change in behavior. The conjugate prior 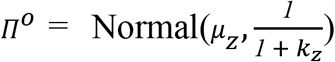 where *μ_Z_* and 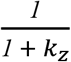 are the mean and variance respectively. Before each trial t, C-TS calculates hypothetical RT *h^t^* directly using the prior hyperparameters (15) and based on the *h^t^* a stimulation site k is selected. Once RT is observed the hyperparameters are updated using posterior update 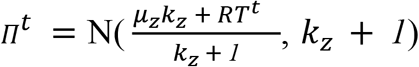 in combination with (10) as shown in Table 1[51].

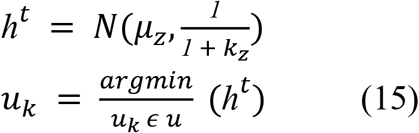

#### Improper priors

As in most MAB algorithms, initially the priors are set as improper priors. To make the posterior proper, we force selection of each stimulation site at least once before control of the selection is given to the optimization algorithm[49].

#### Brute force

We compared all of the above optimizers against simple brute force (BF). The brute force algorithm selects each stimulation site repeatedly for a proportional fraction of the simulated trials. The exact number of applications is determined by the block size, see “Block-wise vs. trial-wise optimization” below. It then declares the optimal site to be the site with the lowest mean RT, i.e the one with maximum stimulation effect.

### Optimizer test with simulated stimulation response surfaces

The performance of a given optimizer depends on the problem structure. For instance, if there is a single obvious global best setting, the greedy and brute force algorithms will suffice. If the response surface is shallow, with local minima very close to the global minimum, the more complex optimizers are more likely necessary. Greedy and brute force algorithms also are expected to scale less well as the number of options (stimulation sites) expands, e.g. as DBS leads move from legacy quadripolar designs to modern multi-contact segmented designs[52,53].

We thus simulated a range of increasingly complex optimization problems, from 2 to 8 stimulation sites (Table 2). We modeled the effects of individual sites after the empirical effects observed in [37]. We assumed that all sites either decrease RT (the desired effect) or have no effect. Table 2 gives the values for *b_in_* ‘s effect on *x_base_*; the effect on *x_conflict_* was assumed to be 10-fold smaller (consistent with the empirical findings that VCVS stimulation mainly affects *x_base_*). For each re-simulation of the problem (see below), the order of these sites was randomly shuffled.

**Table 2:** Stimulation sites and stimulation effect for different number of stimulation sites. The units are effectively log-seconds, consistent with the definition of *x_base_*.

| Problem size | Stimulation effect |  |  |  |  |  |  |  |
| --- | --- | --- | --- | --- | --- | --- | --- | --- |
|  | 1 | 2 | 3 | 4 | 5 | 6 | 7 | 8 |
| 8 | 0 | -0.005 | -0.01 | -0.02 | -0.03 | -0.031 | -0.04 | -0.07 |
| 6 | 0 | -0.005 | -0.01 | -0.02 | -0.04 | -0.07 |  |  |
| 4 | 0 | -0.005 | -0.01 | -0.04 |  |  |  |  |
| 2 | -0.01 | -0.04 |  |  |  |  |  |  |

Another key performance factor for an optimization algorithm is its ability to distinguish between two closely spaced minima. We therefore also simulated the 2-site case, but with increasingly small separation between the two options, all the way to a degenerate case with no difference between options. These values b1-b7 are given in Table 3.

**Table 3:** Stimulation effects for 2 stimulation site locations for the optimization problems, with decreasing differences between the 2 sites. *b_ss_* is a special case used only for demonstrating steady-state response/settling of the sensor model. It is specifically chosen to have very large and visibly different effects.

| Stimulation sites | Stimulation effect |  | Stimulation sites | Stimulation effect |  |
| --- | --- | --- | --- | --- | --- |
|  | 1 | 2 |  | 1 | 2 |
| b1 | 0.05 | -0.05 | b5 | 0 | -0.02 |
| b2 | 0 | -0.10 | b6 | 0 | -0.01 |
| b3 | 0 | -0.05 | b7 | 0 | 0 |
| b4 | 0 | -0.03 | $b_{ss}$ | 0.25 | -0.25 |

The primary metric for optimizer evaluation is accuracy, defined as the fraction of times that a given algorithm identifies the known best site.

### Optimization simulations

Optimizer accuracy may be sensitive to the steady-state settling behavior of the specific problem, the convergence rules, and aspects of the problem. We tested the above algorithms across a range of assumptions (Table 4), including:

**Table 4:** Simulation configuration for overall closed loop simulations to identify the effects of different parameters on the algorithm’s accuracy.

|  | Replicates | Trials per replicate | Blocksize | Stimulation sites |
| --- | --- | --- | --- | --- |
| Effect of block-wise vs. trial-wise stimulation | 1000 | 600 | 1,5,10,15,20,25,30,35,40,45,50 | 8 |
| Effect of convergence criteria | 1000 | 100,200,300,400,500,600,700,800,900,1000 | 15 | 8 |
| Effect of number of stimulation sites | 1000 | 600 | 15 | 2,4,6,8 |
| Effect of signal to noise ratio | 1000 | 600 | 15 | 8 |
| Discriminability of different stimulation effects | 1000 | 600 | 15 | 2 |
| Ensemble (1-7) | 1000 | 600 | 15 | 2,4,6,8 |

#### Effect of Block-wise vs. trial-wise stimulation

By default, all the above optimization algorithms explore a new setting (stimulation site) on every trial. This will work poorly, however, with the specific linear dynamical system model underlying our generator-sensor model. As described in [37], the model is meant to smooth out stochastic variation in RTs, and as such, the underlying state variables change slowly in response to perturbation. The effect of a new stimulation site may not be clearly detectable until it has been applied for several trials in a row. We thus compared trial-wise optimization to a block-wise optimization, where the stimulation was only able to change every *n* trials, and where the mean RT over that block of trials is used as the outcome to update the algorithm’s internal model. We refer to *n* as the “blocksize”. The block approach creates a trade-off – higher blocksize will stabilize the estimate and make subtle effects more obvious, but will require more total trials to converge. We thus evaluated convergence properties for blocksize ranging from 1 to 50, with a step size of 5. For each of these, we simulated only 600 trials. This represents 2-3 hours of total optimization time (assuming some breaks), which is about the most a patient can be expected to tolerate in a clinical setting[15]. Using this more limited trial count also helps highlight the trade off just described. For each value of blocksize and for each proposed algorithm, we simulated 1000 task runs each with 600 trials, with a generator randomly drawn from the above generator model pool (1000 models seeded from 6 participants). The sensor model was similarly selected randomly as described under “Sensor model”. For each run, the optimizer was only permitted to choose stimulation sites once all sites had been selected at least once (i.e., we initialized the algorithm by a forced exploration). This leads to prior propers and encourages convergence of the sensor to the true current value of *x_base_* regardless of the value of *x_0_*. We simulated 8 stimulation sites for this problem, as described in Table 2.

#### Effect of convergence criterion

The convergence of a bandit algorithm depends on the complexity of the problem under consideration. Waiting for algorithms to converge could be impractical if the algorithm takes longer to converge, as real humans will have difficulty tolerating more than 2-3 hours of testing. We thus tested how optimization performance depends on the number of trials (a fixed stopping criterion). We tested 100, 200, 300, 400, 500, 600, 700, 800, 900, and 1000 trial runs. The overall simulation was repeated 1000 times with a problem of 8 stimulation sites (Table 2) and blocksize 15. The position of the optimal stimulation site was shuffled for each repetition. The generator models were randomly selected for each simulation run as above.

#### Effect of number of stimulation sites

As the number of available options (stimulation sites) increases, algorithms may take longer to reliably estimate each site’s effect. This could give an advantage to the Bayesian algorithms, which generally explore more efficiently. To test this possibility, we evaluated how algorithm performance depends on the number of potential stimulation sites. For this simulation, we tested 2, 4, 6, and 8 sites, with surfaces described in Table 2. These simulations were repeated for 1000 replicates, with each replicate having 600 trials and a blocksize of 15. The location of the optimal stimulation site was shuffled for each repetition.

#### Effect of state noise/signal to noise ratio

The signal to noise ratio (SNR) influences how well bandit algorithms perform. A noisy system would produce noisy RTs, and it would be difficult for the algorithm to identify any changes caused by the stimulation effect. We evaluated how different noise floors affect the algorithm accuracy. For this simulation, rather than using the values from the grid search, we selected *W* (*σ_1_* and *σ_2_*) as described in Table 5. These values were perturbed by N(0, 0.0001) in order to use a slightly different generator model in each simulation. These values correspond to different SNRs, expressed as the ratio of the largest stimulation effect in b to the size of the noise variance. We simulated each SNR for 1000 replicates, using 600 trials per replicate, a blocksize of 15, and 8 stimulation sites as described in Table 2.

**Table 5:**
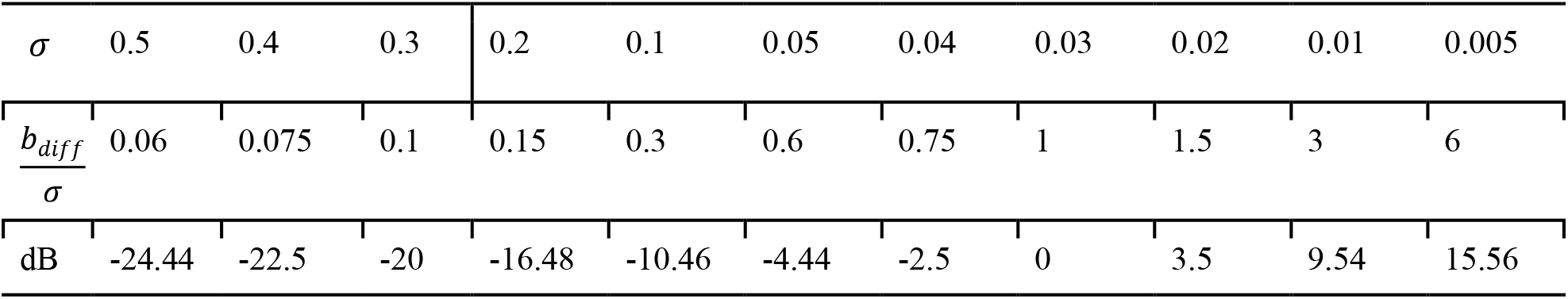
Different state noise levels. *b_diff_* represents the difference between the most effective and second most effective stimulation site.

#### Discriminability of different stimulation effects

It may be challenging to detect a global optimum if two sites have very similar performance. We evaluated how close two stimulation sites could be while still being reliably distinguishable. For simulations, generators were randomly selected from the generator pool and sensor models selected as described in “Sensor model”. Each row of Table 3 was simulated with 1000 replicates, 600 trials per replicate, 2 stimulation sites, and blocksize of 15.

### Ensemble simulations

Even the best optimization algorithm can still converge away from the global optimum, particularly if the measurement (RT) is noisy. This might be mitigated by an ensemble approach, where optimization is performed repeatedly from different initial conditions, or simply by allowing the random choices inherent in the algorithms to proceed from different seeds. Clinically, this could be realized by redoing the optimization with the same patient on multiple days, as has been suggested in [34]. We simulated this ensemble case, by repeatedly re-simulating the same generator and different sensor model. We tested ensembles of 1, 2, 3, 4, 5, 6 and 7 repetitions over 1000 generator-sensor pairs selected as described above, with 600 trials per replicate, blocksize 15, stimulation problem of size 2, 4, 6 and 8, and noise level as defined in the “Grid search for model noise processes” section. The selection for computing accuracy was then based on a hard majority vote across repetitions.

## Results

Successful optimization depends on: 1) selection of reasonable parameters for the sensor model, particularly the noise process terms, 2) the generator’s ability to simulate RTs with a distribution comparable to actual participants, 3) the sensor model’s ability to accurately track the generator’s underlying unobservable states, without direct information about them and 4) the selection of an optimizer that can reliably converge to the correct answer in this specific use case.

### Model noise process determination via grid search

Modeling includes estimation of parameters using the COMPASS EM algorithm. The full parameter estimation problem given in (3) has many unknowns, which in practice often leads to convergence on a local maximum. This is particularly problematic when estimating the state noise *W*, leading to the simplifications in (4) and (5). To enable those simplifications, we performed a grid search to identify reasonable fixed/cross-patient values for *W*, seeking values that minimize the deviance between actual and predicted data during the expectation step of EM.

Figure 2 shows the deviance for the state-space model of (1) and (2), fitted using different noise variances for *x_base_* and *x_conflict_*. A region of values corresponding to [4,19] (*σ*_1_ ≡ [0.0014,0.25]) for *x_base_* and [2,34] (*σ*_2_ ≡ [0.000007629,0.5]) for *x_conflict_* produced small deviance on average (Figure 2A). Due to the unavailability of ground truth about optimal scale values, the specific best point within this region cannot be determined. In practice, any *W* drawn from within the low-deviance zones would likely work well. We verified that this region of low deviance is present in single-participant data (examples in Figure 2B-C). In inspecting those data, we noted that some participants (e.g., S1 shown in Figure 2B) have a narrower range of acceptable *W* values. We thus used this narrower range for all further simulations described below. This corresponded to scale [7,16] (*σ*_1_ ≡ [0.0039, 0.0884]) for *x_base_* and [13,27] (*σ*_2_ ≡ [0.00008,0.0110]) for *x_conflict_*.

**Figure 2:**
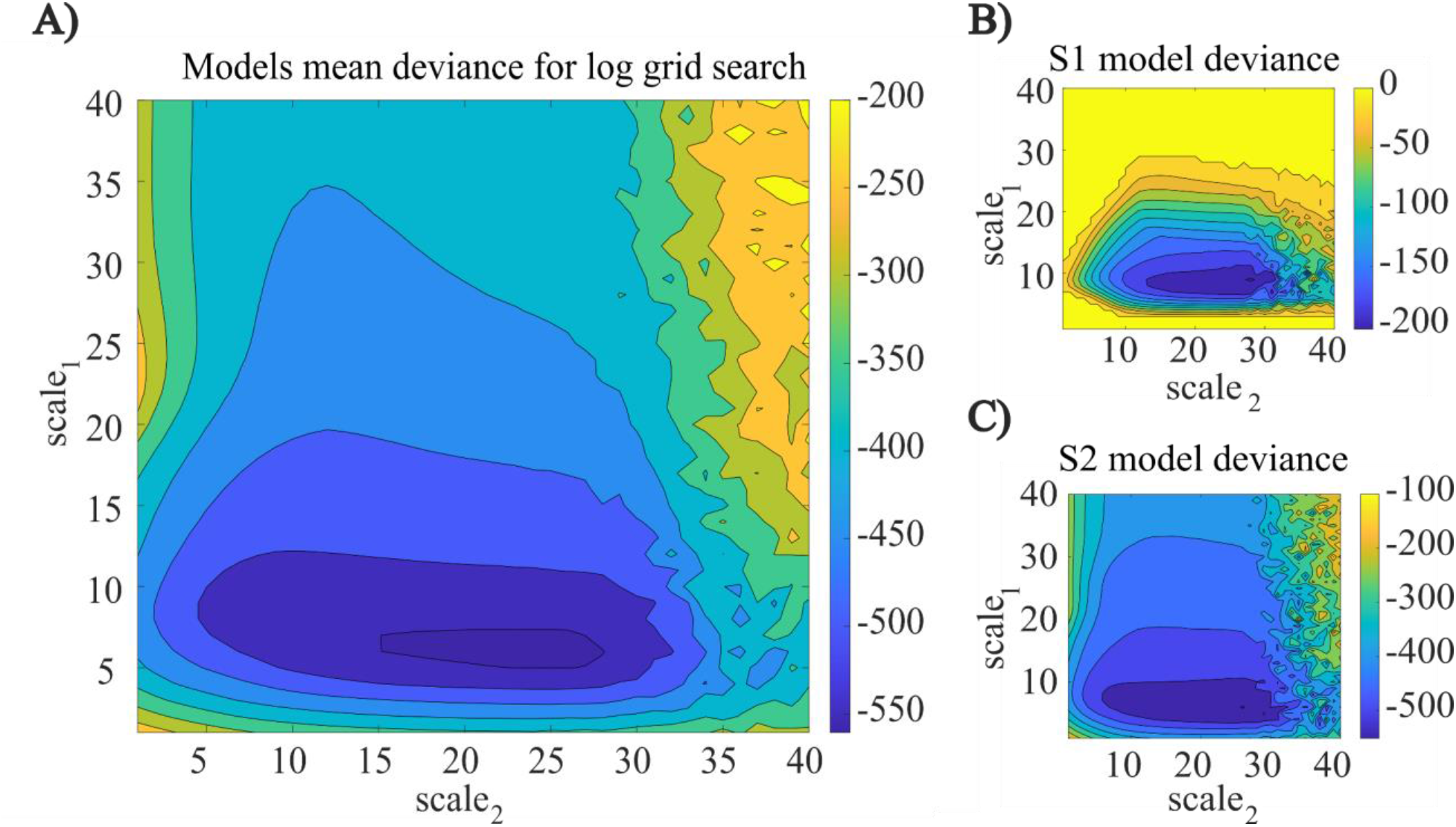
Logarithmic grid search for *W*. The deviance is calculated for the model fitted with different values of *σ_1_* and *σ_2_* (here expressed as scale factors, see equation 6). Deviance values greater than 0 were replaced by 0 to better highlight the contours. A) Averaged deviance across individuals. There is a zone of low deviance across participants. B-C) Examples of the same deviance plot from 1000 simulations of individual participants. The region shown in (A) is present in the individual plots, i.e. is not an artifact of averaging.

### Generator Model Validation

A good model fit using (4) and (5) acts as an approximation of actual participants receiving stimulation, which means that a good model would generate the same distribution as that of the participants’ (S1-S6) data. However, empirically due to the assumptions for (4) and (5), the estimated model requires additional model tuning for exactly matching the distribution of any individual participant. We validated that the generator models, with and without tuning, produced plausible behavior.

Without any tuning (simple fitting to empirical data via COMPASS), the RTs emitted by the generator follow the expected gamma distribution (Figure 3A). While not a good model for S1 specifically, the generator model output thus follows the expected statistics of actual patient data, i.e. could be used as an example of a new, unobserved patient. To verify this, we fit a gamma distribution to the simulated RT, then calculated the empirical distribution of the RTs expected given that fit. The KS distance between the generator-model RT’s probability distribution function and the probability distribution function of a gamma distribution corresponded to p>0.05 in 100% of the simulation runs across patients. As further verification, we showed that interference trials have higher RTs by the amount that would be expected during MSIT (Figure 3B). We also verified that, if necessary, a specific patient’s data could be matched through the generator model. Hand-tuning parameters A, *μ*, w, v and *α* generated simulations that almost perfectly matched with the empirical observations (Figure 3A).

**Figure 3:**
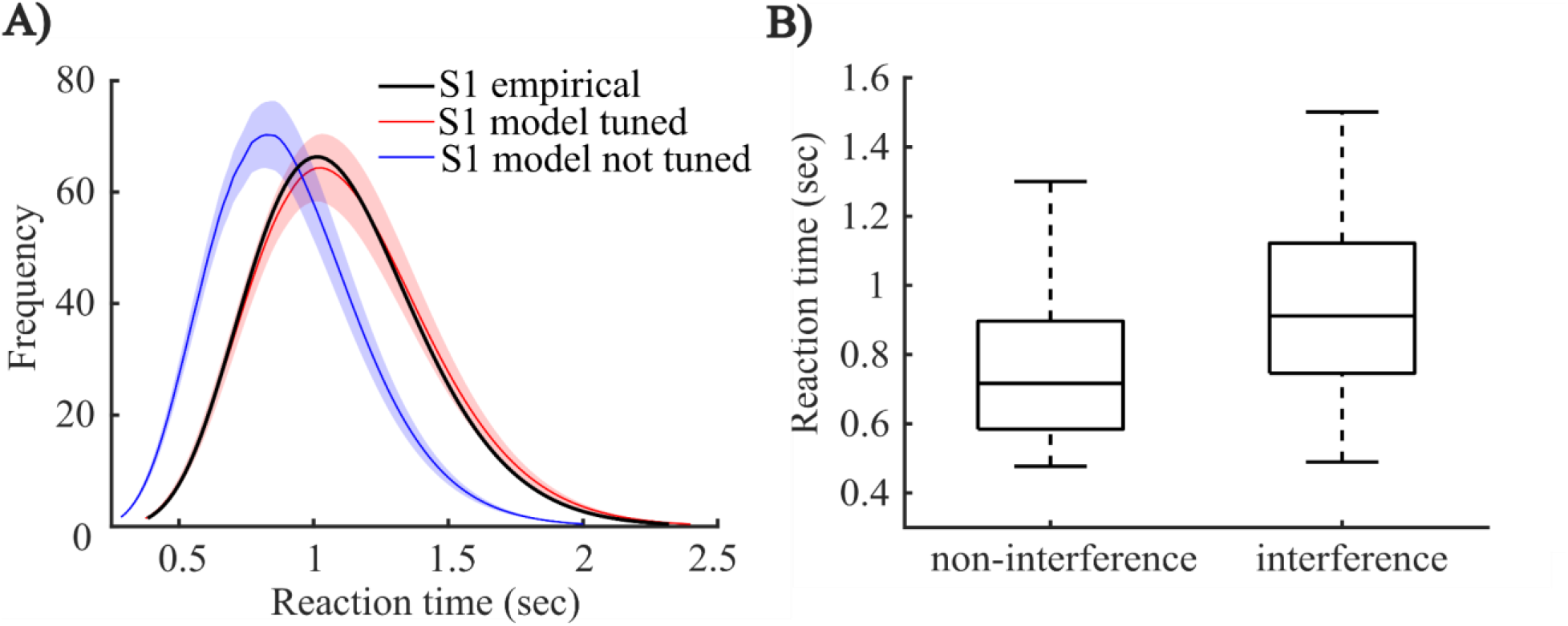
Simulations of MSIT behavior as emitted by the generator model. A) Comparison of actual and simulated behavior (RT distributions) for S1. The curves represent the mean distribution of 1000 runs of 1000 simulated trials each, with trials randomly set as interference/non-interference. The shaded bound represents the standard error of that mean over 1000 re-simulations. The empirical behavior (black) matches well with the RT distribution emitted by a tuned model fit to those data (red). The distribution of an untuned model has the same overall shape, but a different mode and slightly different skew, demonstrating that slight changes to the model parameters can effectively simulate new patients. B) Distribution of RTs during interference and non-interference conditions for 1000 simulated trials from S2. As expected, interference trials have higher RT by approximately 200 ms.

### Sensor model tracking of ground truth and steady state response

The sensor model tracks the generator well (Figure 4A-C) across the simulations, with a median NRMSE of 0.0935 for the actual/estimated RT, 0.0541 for *x_base_*, and 0.0784 for *x_conflict_*. This suggests that the sensor models can accurately track the unobservable generator state, and that this holds true across many random realizations. However, this tracking is not instantaneous. As seen particularly in Figure 4C, the nature of the sensor model, the random initialization of *x_0_*, and the chosen noise covariances cause the tracking to be temporarily inaccurate for the first 20 trials.

**Figure 4:**
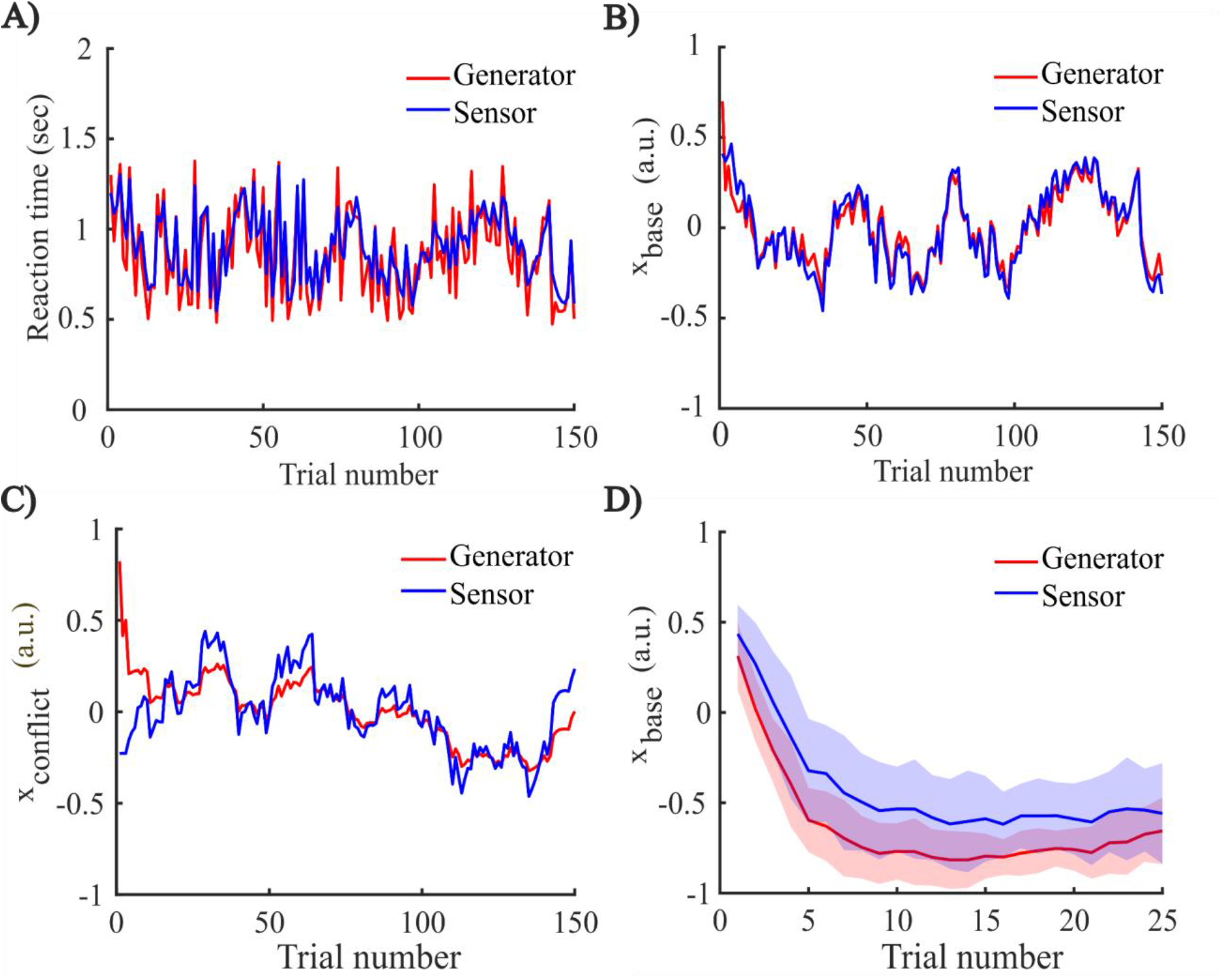
Sensor model tracking the generator’s output and unobserved internal states. A) Comparison between emitted generator RTs and sensor’s prediction of those RTs. This panel shows the first 150 out of 1000 simulated trials. As expected, the sensor output is a smoothed (expected value) version of the generator output. B-C), sensor state estimates compared to generator ground truth. There is accurate tracking of both states after about 20 trials, with this “lead in” period necessary because *x_0_* of the sensor and generator models are initialized to different random numbers. D) Generator and sensor steady state response for a single patient. At trial 0, stimulation switches between the two values of *b_ss_*, applying a strong forcing. The generator’s internal state settles to steady state in roughly 10 trials, and the sensor tracks this closely. Line represents the mean value of *x_base_* for generator and sensor over a blocksize of 25 with 40 replicates (1000 trials), whereas the shading represents the standard error of the mean over replicates.

Similarly, the model structure means that the tracked state variable *x_base_* does not change immediately when stimulation is applied. Since we assume a linear dynamical system with a substantial autoregressive component, it takes 5-10 trials for the effect of a continuous perturbation to settle to a new steady state (Figure 4D). In general, the number of trials to reach steady state is dependent on the stimulation effect strength, i.e. stronger perturbations will take longer, but also will be more easily detected despite small stochastic variations. Regardless, this result argues that the effects of stimulation can only be evaluated over blocks of trials, not on individual trials.

### Bayesian Optimization

Having validated the generator and sensor model and identified settling time as a consideration, we evaluated which optimization algorithms could best identify optimal stimulation sites for improving MSIT performance (decreasing task RTs). We measured the accuracy (probability of identifying a known global optimum) of multiple algorithms as a function of 1) Blocksize for stimulation, 2) Number of stimulation sites (“problem size”), 3) Number of trials performed, 4) Distance between the stimulation effects of the stimulation sites, 5) Signal to noise ratio of stimulation effect relative to state noise, and 6) Number of times an algorithm is repeated during an ensemble strategy.

### Blocksize for stimulation

The accuracy of all algorithms improves with increasing blocksize, but plateaus around a block size of 10 (Figure 5A and Table 6). This is visible across all algorithms. This is consistent with the observation in Figure 4 that the stimulation effect is not instantaneous, but takes a few trials to manifest in *x_base_*. Further, at these higher blocksizes, UCB1 is generally the best algorithm. When we compared all algorithms across problem sizes and blocksizes, UCB1 was the most likely to converge to the known optimum (Table 6). Bayesian algorithms were the best for small blocksizes and single trial simulations, but lost this advantage in part because as the blocksize increases, the estimate of *x_base_* tends to the actual *x_base_*. At the same time, most algorithms consistently outperform brute force, emphasizing the overall value of more advanced optimization strategies.

**Figure 5:**
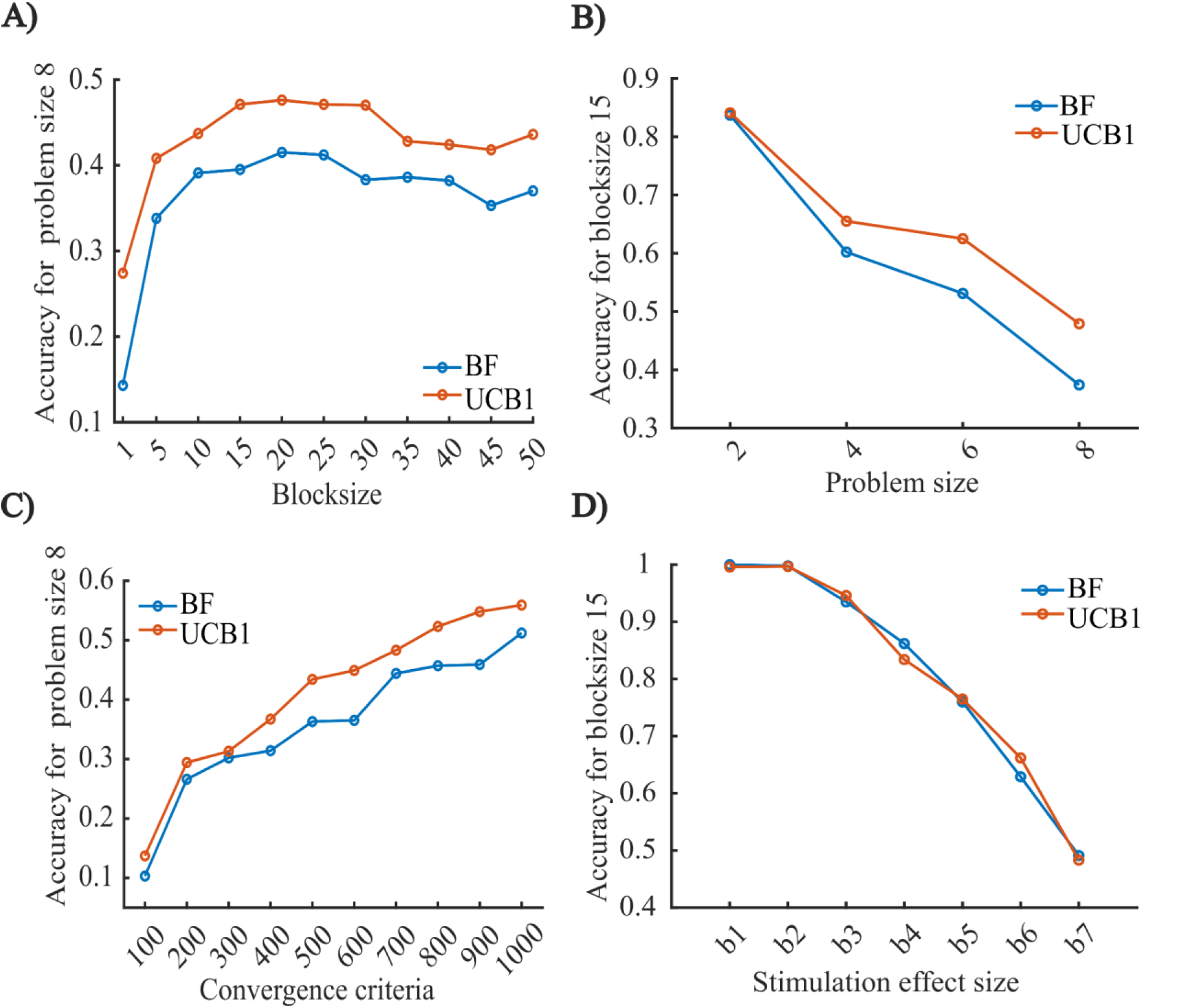
Comparative performance of brute force (BF) and UCB1 algorithms, all from 1000 replicates, with block, problem structure, and number of trials varied as per Methods. A) Performance for a fixed optimization problem (8 sites), at increasing blocksize. Performance plateaus at blocksize of 10 and above. For simplicity, we show UCB1 as the overall best performer and BF as the simplest/default comparison; see Table 6 for detailed results on all algorithms. B) comparative performance of UCB1 with BF for increasing numbers of stimulation sites. Both algorithms lose performance as the problem becomes more complex, but UCB1 consistently performs better on harder problems. C) Improved performance with more trials to convergence. If the number of trials is increased, specifically above the 600 trials shown in A-B, accuracy also increases (see Table 7). D) resolution of closely spaced minima. As the stimulation effect of both the sites approaches the same value, the ability of the algorithms to identify the optimal stimulation site decreases. However, performance continues to exceed chance until the degenerate case of b7 (no difference).

**Table 6:** Different algorithm performance with different blocksize for problem sizes of 4, 6 and 8 sites. The bold values show algorithm accuracies that exceed brute force (BF). Red-colored values show the best algorithm for the given blocksize. The final column, “Wins”, indicates the number of times that a given algorithm was best across all blocksizes. For all problem sizes, UCB1 had the most wins, i.e. it was more likely than other algorithms to find the best solution.

Blocksize for stimulation
| Problem size 4 |  | Blocksize |  |  |  |  |  |  |  |  |  | Wins |
| --- | --- | --- | --- | --- | --- | --- | --- | --- | --- | --- | --- | --- |
| Algorithms | 1 | 5 | 10 | 15 | 20 | 25 | 30 | 35 | 40 | 45 | 50 |  |
| BF | 0.307 | 0.47 | 0.602 | 0.615 | 0.642 | 0.614 | 0.625 | 0.644 | **0.66** | **0.637** | 0.61 | 2 |
| Greedy | 0.28 | 0.392 | 0.538 | 0.582 | 0.576 | 0.598 | 0.638 | 0.602 | 0.612 | 0.615 | 0.651 | 0 |
| ε-greedy | **0.398** | **0.522** | 0.554 | 0.586 | 0.603 | 0.612 | 0.637 | 0.619 | 0.63 | 0.616 | 0.637 | 0 |
| UCB1 | **0.439** | **0.589** | **0.675** | **0.664** | **0.679** | **0.637** | **0.67** | **0.677** | 0.654 | 0.628 | **0.679** | 6 |
| Bayes-UCB | **0.475** | **0.573** | **0.631** | **0.618** | **0.652** | **0.662** | **0.649** | **0.663** | 0.631 | 0.629 | **0.636** | 2 |
| TS-Bernoulli | **0.417** | **0.542** | **0.639** | **0.619** | 0.612 | **0.635** | 0.593 | 0.603 | 0.61 | 0.582 | **0.641** | 0 |
| TS-Poisson | **0.459** | **0.53** | 0.591 | 0.586 | 0.603 | **0.643** | 0.612 | 0.633 | 0.619 | 0.626 | **0.647** | 0 |
| TS-Normal | **0.4** | **0.601** | **0.612** | **0.623** | 0.619 | **0.618** | **0.631** | 0.623 | 0.611 | 0.595 | 0.594 | 1 |
| C-TS | **0.356** | **0.527** | 0.522 | 0.553 | 0.582 | 0.555 | 0.556 | 0.55 | 0.593 | 0.583 | 0.588 | 0 |

| Problem size 6 |  | Blocksize |  |  |  |  |  |  |  |  |  | Wins |
| --- | --- | --- | --- | --- | --- | --- | --- | --- | --- | --- | --- | --- |
| Algorithms | 1 | 5 | 10 | 15 | 20 | 25 | 30 | 35 | 40 | 45 | 50 |  |
| BF | 0.213 | 0.447 | 0.526 | 0.511 | 0.546 | 0.561 | 0.568 | 0.547 | 0.556 | 0.533 | 0.547 | 0 |
| Greedy | **0.216** | 0.36 | **0.523** | **0.597** | **0.561** | **0.566** | **0.592** | **0.569** | **0.597** | **0.569** | **0.585** | 3 |
| ε-greedy | **0.367** | **0.465** | **0.542** | **0.554** | **0.603** | **0.599** | **0.57** | **0.587** | **0.591** | **0.564** | **0.581** | 0 |
| UCB1 | **0.383** | **0.508** | **0.572** | **0.568** | **0.616** | **0.586** | **0.618** | **0.601** | **0.585** | **0.6** | **0.564** | 5 |
| Bayes-UCB | **0.428** | **0.55** | **0.567** | **0.569** | **0.588** | **0.584** | **0.585** | **0.556** | **0.573** | **0.563** | 0.541 | 2 |
| TS-Bernoulli | **0.358** | **0.544** | **0.551** | **0.542** | **0.557** | **0.536** | 0.546 | **0.549** | 0.501 | **0.549** | 0.516 | 0 |
| TS-Poisson | **0.408** | **0.527** | **0.533** | **0.566** | **0.566** | **0.61** | **0.589** | 0.542 | **0.578** | **0.561** | **0.554** | 1 |
| TS-Normal | **0.316** | **0.522** | **0.534** | **0.567** | **0.553** | **0.56** | 0.562 | **0.554** | 0.494 | 0.495 | 0.537 | 0 |
| C-TS | **0.281** | 0.435 | 0.449 | 0.483 | 0.496 | 0.497 | 0.517 | 0.529 | 0.552 | 0.514 | **0.551** | 0 |

Table 6: Different algorithm performance with different blocksize for problem sizes of 4, 6 and 8 sites.
| Problem size 8 |  | Blocksize |  |  |  |  |  |  |  |  |  | Wins |
| --- | --- | --- | --- | --- | --- | --- | --- | --- | --- | --- | --- | --- |
| Algorithms | 1 | 5 | 10 | 15 | 20 | 25 | 30 | 35 | 40 | 45 | 50 |  |
| BF | 0.163 | 0.334 | 0.4 | 0.388 | 0.403 | 0.4 | 0.39 | 0.387 | 0.363 | 0.379 | 0.362 | 0 |
| Greedy | 0.162 | 0.33 | **0.419** | **0.457** | **0.455** | **0.439** | **0.467** | **0.42** | **0.454** | **0.407** | **0.427** | 1 |
| ε-greedy | **0.279** | **0.387** | **0.443** | **0.441** | **0.42** | **0.449** | **0.444** | **0.437** | **0.474** | **0.439** | **0.408** | 3 |
| UCB1 | **0.24** | **0.37** | **0.456** | **0.5** | **0.472** | **0.456** | **0.437** | **0.41** | **0.429** | **0.423** | **0.439** | 5 |
| Bayes-UCB | **0.343** | **0.394** | **0.412** | **0.406** | **0.423** | 0.394 | 0.382 | 0.353 | **0.418** | **0.4** | **0.367** | 2 |
| TS-Bernoulli | **0.267** | **0.368** | 0.384 | **0.403** | 0.383 | 0.379 | 0.377 | **0.405** | **0.374** | 0.374 | **0.382** | 0 |
| TS-Poisson | **0.322** | **0.372** | **0.416** | **0.434** | **0.411** | 0.391 | 0.388 | 0.385 | **0.381** | 0.35 | **0.389** | 0 |
| TS-Normal | **0.204** | **0.368** | **0.4** | **0.406** | **0.405** | **0.409** | 0.368 | 0.348 | 0.352 | 0.367 | **0.372** | 0 |
| C-TS | **0.206** | 0.278 | 0.299 | 0.357 | 0.378 | 0.397 | 0.384 | **0.388** | **0.395** | **0.393** | **0.401** | 0 |

### Problem size

Accuracy of all algorithms decreases with increases in the number of stimulation sites. This is a consequence of a decrease in the available samples for a fixed convergence criterion (number of trials) and blocksize. All algorithms outperformed brute force more strongly as the problem size (number of sites) increased (Figure 5B). However, UCB1 maintained its advantage over the Bayesian and greedy algorithms (Table 6). The decrease in accuracy affects the Bayesian algorithms more since they are based on inferring the underlying distribution of the data, whereas greedy algorithms comparatively start performing better.

### Convergence criterion

Accuracy increases with increase in the convergence criterion (number of allowed trials). However, in practical situations the number of trials is limited, hence our emphasis on 600 trials across other simulations. In this analysis UCB1 again outperformed other algorithms (Table 7). It specifically maintained its advantage over brute force at all convergence criteria above 500 trials (Figure 5C), again demonstrating the benefits of a more efficiently exploring optimizer.

**Table 7:** Different algorithm performance with different convergence criteria. UCB1 outperforms all other algorithms across problem sizes (most Wins). Convergence and accuracy are linearly related.

| Problem size 4 |  | Convergence Criteria |  |  |  |  |  |  |  |  |  |
| --- | --- | --- | --- | --- | --- | --- | --- | --- | --- | --- | --- |
| Algorithms | 100 | 200 | 300 | 400 | 500 | 600 | 700 | 800 | 900 | 1000 | Wins |
| BF | 0.36 | 0.438 | 0.518 | 0.548 | 0.577 | 0.615 | 0.626 | 0.692 | 0.687 | 0.7 | 0 |
| Greedy | **0.388** | **0.484** | 0.5 | **0.556** | 0.532 | 0.563 | 0.573 | 0.576 | 0.587 | 0.597 | 1 |
| ε-greedy | 0.355 | **0.444** | 0.495 | **0.562** | **0.581** | 0.608 | 0.607 | 0.614 | 0.647 | 0.649 | 0 |
| UCB1 | **0.402** | **0.467** | **0.54** | **0.57** | **0.628** | **0.66** | **0.664** | **0.701** | **0.705** | **0.751** | 9 |
| Bayes-UCB | **0.373** | **0.453** | 0.502 | **0.568** | **0.609** | 0.608 | **0.65** | 0.691 | **0.72** | **0.723** | 0 |
| TS-Bernoulli | **0.362** | **0.439** | 0.485 | 0.527 | **0.589** | 0.607 | **0.629** | 0.673 | 0.655 | 0.694 | 0 |
| TS-Poisson | 0.344 | **0.474** | 0.511 | 0.537 | **0.579** | **0.619** | **0.631** | 0.629 | 0.646 | 0.695 | 0 |
| TS-Normal | **0.362** | **0.454** | 0.487 | 0.532 | **0.584** | **0.657** | **0.653** | 0.675 | **0.703** | **0.713** | 0 |
| C-TS | 0.35 | 0.431 | 0.48 | 0.512 | 0.523 | 0.542 | 0.589 | 0.586 | 0.595 | 0.621 | 0 |

| Problem size 6 |  | Convergence Criteria |  |  |  |  |  |  |  |  | Wins |
| --- | --- | --- | --- | --- | --- | --- | --- | --- | --- | --- | --- |
| Algorithms | 100 | 200 | 300 | 400 | 500 | 600 | 700 | 800 | 900 | 1000 |  |
| BF | **0.313** | 0.357 | 0.408 | 0.501 | 0.527 | 0.534 | 0.595 | 0.623 | 0.634 | 0.635 | 1 |
| Greedy | 0.293 | **0.401** | **0.45** | **0.507** | **0.535** | **0.584** | 0.561 | 0.576 | 0.59 | 0.6 | 0 |
| ε-greedy | 0.28 | **0.427** | **0.47** | **0.505** | **0.545** | **0.537** | 0.569 | 0.62 | 0.611 | **0.647** | 1 |
| UCB1 | 0.284 | **0.421** | **0.474** | **0.559** | **0.544** | **0.62** | **0.615** | **0.661** | **0.675** | **0.711** | 7 |
| Bayes-UCB | 0.291 | **0.376** | **0.418** | 0.479 | 0.515 | **0.56** | 0.586 | 0.619 | **0.667** | **0.667** | 0 |
| TS-Bernoulli | 0.282 | 0.344 | 0.393 | 0.477 | 0.492 | **0.571** | 0.56 | 0.601 | 0.615 | **0.653** | 0 |
| TS-Poisson | 0.304 | **0.39** | **0.424** | 0.446 | **0.537** | **0.546** | 0.592 | 0.594 | **0.641** | **0.666** | 0 |
| TS-Normal | 0.284 | **0.376** | **0.431** | 0.484 | **0.529** | **0.551** | **0.603** | 0.611 | **0.638** | **0.665** | 0 |
| C-TS | 0.287 | **0.37** | 0.388 | 0.409 | 0.446 | 0.459 | 0.503 | 0.513 | 0.517 | 0.557 | 0 |

| Problem size 8 |  | Convergence Criteria |  |  |  |  |  |  |  |  | Wins |
| --- | --- | --- | --- | --- | --- | --- | --- | --- | --- | --- | --- |
| Algorithms | 100 | 200 | 300 | 400 | 500 | 600 | 700 | 800 | 900 | 1000 |  |
| BF | 0.103 | 0.266 | 0.302 | 0.314 | 0.363 | 0.365 | 0.444 | 0.457 | 0.459 | 0.512 | 0 |
| Greedy | **0.12** | **0.269** | **0.357** | **0.395** | **0.405** | **0.481** | **0.45** | **0.477** | **0.477** | 0.508 | 3 |
| ε-greedy | **0.132** | **0.266** | **0.327** | **0.395** | **0.454** | **0.425** | **0.467** | **0.507** | **0.515** | **0.524** | 2 |
| UCB1 | **0.137** | **0.294** | **0.313** | **0.367** | **0.434** | **0.449** | **0.483** | **0.523** | **0.548** | **0.559** | 5 |
| Bayes-UCB | **0.138** | 0.247 | 0.281 | **0.321** | **0.379** | **0.402** | 0.417 | **0.499** | **0.508** | 0.493 | 1 |
| TS-Bernoulli | **0.12** | 0.245 | **0.303** | **0.322** | **0.365** | **0.387** | 0.413 | **0.458** | 0.456 | 0.473 | 0 |
| TS-Poisson | **0.107** | **0.275** | 0.27 | **0.317** | **0.363** | **0.384** | 0.423 | **0.459** | **0.507** | **0.542** | 0 |
| TS-Normal | **0.135** | 0.244 | 0.274 | **0.339** | 0.355 | **0.404** | 0.436 | **0.475** | **0.493** | **0.541** | 0 |
| C-TS | **0.104** | 0.228 | 0.292 | **0.322** | 0.311 | 0.331 | 0.348 | 0.38 | 0.378 | 0.396 | 0 |

### Discriminability of different stimulation effects

Accuracy depends on the closeness of local optima to the global optimum. When given two stimulation choices to optimize, brute force and UCB1 could identify the optimum with probability above chance, even in very hard problems (Figure 5D). They only fell to chance in the degenerate case b7, when the two sites were identical. In this specific analysis, there was no advantage for UCB1 over BF; both had almost identical performance as discriminability fell.

### Signal to noise ratio

Accuracy of the algorithms is sensitive to noise. In other words, if there is more noise in the data generator, it will mask the stimulation effect and the overall accuracy will decrease. We compared algorithm performance across different levels of SNR, defined based on ratio of the stimulation effect to the state noise process variance. UCB1’s advantage was less clear in this case; Bayesian and even greedy algorithms could outperform it in specific cases, although UCB1 was still either first or second across problem sizes (Table 8).

**Table 8:** Algorithm performance across varying SNRs. UCB1 outperformed or tied all other algorithms at problem sizes 4 and 6, whereas *∊*-greedy performed better for the most challenging problem of 8 sites.

| Problem size 4 |  | SNR(dB) |  |  |  |  |  |  |  |  |  |  |
| --- | --- | --- | --- | --- | --- | --- | --- | --- | --- | --- | --- | --- |
| Algorithms | -24.44 | -22.5 | -20 | -16.48 | -10.46 | -4.44 | -2.5 | 0 | 3.52 | 9.54 | 15.56 | Wins |
| BF | 0.372 | 0.381 | 0.394 | 0.468 | 0.549 | 0.602 | 0.619 | 0.649 | 0.668 | 0.708 | 0.715 | 0 |
| Greedy | 0.326 | 0.353 | 0.371 | 0.41 | 0.493 | 0.538 | 0.551 | 0.557 | 0.571 | 0.638 | 0.651 | 0 |
| ε-greedy | 0.337 | 0.366 | 0.385 | 0.442 | 0.497 | 0.585 | 0.611 | 0.61 | 0.643 | 0.647 | 0.7 | 0 |
| UCB1 | **0.408** | **0.405** | **0.399** | 0.466 | 0.534 | **0.607** | **0.635** | **0.699** | **0.737** | **0.717** | **0.778** | 4 |
| Bayes-UCB | 0.322 | 0.403 | **0.427** | 0.432 | 0.529 | **0.649** | **0.642** | **0.653** | **0.701** | **0.761** | **0.811** | 4 |
| TS-Bernoulli | 0.342 | 0.368 | **0.401** | 0.438 | **0.557** | 0.56 | 0.583 | **0.679** | 0.665 | **0.728** | **0.77** | 0 |
| TS-Poisson | 0.369 | 0.385 | **0.395** | 0.44 | 0.507 | 0.59 | 0.615 | 0.623 | **0.681** | **0.714** | **0.784** | 0 |
| TS-Normal | 0.362 | 0.37 | **0.401** | **0.469** | **0.549** | **0.607** | 0.616 | **0.652** | **0.682** | **0.763** | **0.807** | 2 |
| C-TS | 0.34 | 0.347 | 0.355 | 0.388 | 0.479 | 0.505 | 0.534 | 0.545 | 0.621 | 0.696 | 0.701 | 0 |

| Problem size 6 |  | SNR(dB) |  |  |  |  |  |  |  |  |  |  |
| --- | --- | --- | --- | --- | --- | --- | --- | --- | --- | --- | --- | --- |
| Algorithms | -24.44 | -22.5 | -20 | -16.48 | -10.46 | -4.44 | -2.5 | 0 | 3.52 | 9.54 | 15.56 | Wins |
| BF | 0.275 | 0.303 | 0.345 | 0.385 | 0.487 | 0.518 | 0.548 | 0.564 | 0.581 | 0.632 | 0.647 | 0 |
| Greedy | 0.268 | 0.299 | **0.345** | **0.395** | **0.491** | **0.543** | 0.545 | **0.574** | **0.612** | 0.625 | **0.649** | 0 |
| ε-greedy | **0.278** | **0.332** | **0.35** | **0.397** | 0.483 | **0.559** | **0.59** | **0.592** | **0.624** | **0.632** | **0.694** | 2 |
| UCB1 | **0.288** | 0.3 | **0.368** | **0.434** | **0.527** | **0.572** | **0.59** | **0.638** | **0.661** | **0.704** | **0.675** | 8 |
| Bayes-UCB | **0.29** | **0.304** | 0.334 | **0.416** | **0.498** | **0.566** | **0.579** | **0.635** | **0.66** | **0.697** | **0.711** | 1 |
| TS-Bernoulli | **0.281** | **0.323** | 0.318 | 0.369 | 0.442 | **0.537** | 0.544 | 0.563 | **0.602** | **0.654** | **0.677** | 0 |
| TS-Poisson | **0.279** | **0.316** | **0.353** | 0.362 | 0.473 | **0.545** | **0.572** | **0.578** | **0.629** | **0.654** | **0.698** | 1 |
| TS-Normal | **0.28** | 0.271 | 0.313 | **0.395** | 0.457 | **0.541** | **0.551** | **0.588** | **0.607** | **0.662** | **0.678** | 0 |
| C-TS | 0.236 | 0.295 | 0.262 | 0.338 | 0.399 | 0.481 | 0.463 | 0.458 | 0.53 | 0.588 | 0.643 | 0 |

Table 8: Algorithm performance across varying SNRs. UCB1 outperformed or tied all other algorithms at problem sizes 4 and 6, whereas ε-greedy performed better for the most challenging problem of 8 sites.
| Problem size 8 |  | SNR(dB) |  |  |  |  |  |  |  |  |  |  |
| --- | --- | --- | --- | --- | --- | --- | --- | --- | --- | --- | --- | --- |
| Algorithms | -24.44 | -22.5 | -20 | -16.48 | -10.46 | -4.44 | -2.5 | 0 | 3.52 | 9.54 | 15.56 | Wins |
| BF | 0.212 | 0.209 | 0.221 | 0.259 | 0.348 | 0.375 | 0.397 | 0.401 | 0.438 | 0.474 | 0.492 | 0 |
| Greedy | **0.221** | **0.236** | **0.255** | **0.303** | **0.387** | **0.451** | **0.458** | **0.476** | **0.52** | **0.533** | **0.548** | 3 |
| ε-greedy | **0.246** | **0.239** | **0.263** | **0.275** | **0.378** | **0.436** | **0.44** | **0.489** | **0.53** | **0.534** | **0.563** | 5 |
| UCB1 | **0.232** | **0.245** | **0.24** | **0.32** | **0.369** | **0.44** | **0.454** | **0.453** | **0.502** | **0.529** | **0.577** | 3 |
| Bayes-UCB | **0.214** | **0.224** | **0.229** | **0.26** | **0.349** | **0.38** | **0.4** | **0.43** | **0.476** | **0.487** | **0.507** | 0 |
| TS-Bernoulli | 0.209 | 0.199 | 0.214 | 0.229 | 0.337 | 0.34 | **0.409** | **0.401** | **0.459** | **0.499** | **0.504** | 0 |
| TS-Poisson | 0.19 | **0.217** | **0.259** | **0.264** | 0.343 | **0.403** | **0.411** | **0.437** | **0.441** | **0.495** | **0.539** | 0 |
| TS-Normal | 0.207 | **0.219** | **0.226** | **0.259** | 0.322 | **0.382** | **0.403** | **0.431** | **0.457** | 0.466 | **0.509** | 0 |
| C-TS | 0.183 | 0.2 | **0.221** | 0.256 | 0.287 | 0.316 | 0.343 | 0.371 | 0.401 | 0.425 | 0.476 | 0 |

### Ensemble method for increased accuracy

Ensemble optimization further improved performance (Figure 6). Even in the most difficult problem of 8 stimulation sites, UCB1 maintained accuracy of 78%, as compared to 47% when simulated once. In simpler problems, e.g. 4 stimulation sites as was originally tested in [37], UCB1’s accuracy exceeded 90%. Performance slightly improved as the ensemble process lengthened from 3 to 5 days, but only asymptotically.

**Figure 6:**
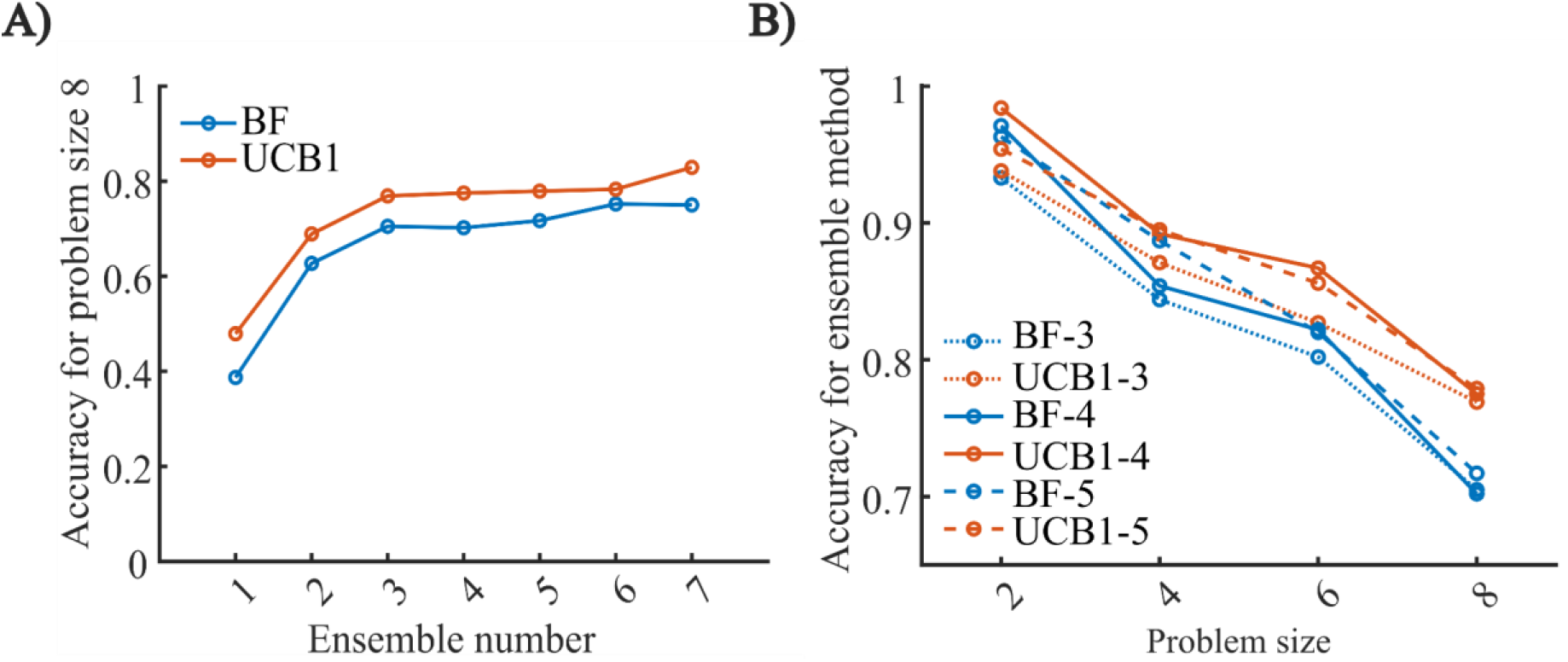
Ensemble algorithm performance. Both UCB1 and BF improve dramatically when optimization is repeatedly run on the same simulated subject. However, UCB1 benefits more, retaining ~80% accuracy even in the most difficult problems. A) Accuracy for increasing number of ensemble runs, showing a plateau after 3 repetitions. B) Accuracy as a function of problem size. UCB1’s advantage over BF increases as the problem becomes more difficult, and this is not mitigated by increasing numbers of ensemble runs. Different numbers (and corresponding linestyles) represent the number of ensemble runs, e.g. UBC1-4 represents 4 repeated UCB1 runs on 4 separate days.

## Discussion

We have demonstrated an online approach to optimizing DBS programming for psychiatric disorders, and specifically, selection of the optimal site during a monopolar survey in the VCVS target. This approach is based on a robust framework for modeling and tracking individual behavior during a cognitive conflict task and evaluating the effect of different DBS contacts/settings on task performance. We have shown that models in this framework can rapidly be fit/adapted to individual participants, and that the most challenging parameter, the state noise covariance, does not need to be patient-specific. Using that general covariance, we have specifically shown that our model framework can track known ground truth, and can detect the effects of stimulation changes within a few task trials. Further, any individual can be precisely replicated and simulated by appropriate model parameter tuning. Finally, we showed that combining this modeling framework with adaptive optimization algorithms is an effective approach to identifying stimulation sites/parameters for a patient. The Upper Confidence Bound (UCB1) algorithm in particular outperforms the brute force approach described in [37] and a variety of alternate algorithms. When deployed in an ensemble, it can reliably identify a known optimum under realistic assumptions of problem size and time available for optimization. As presented above, specific choices within the optimization (block size, number of trials per run, ensemble structure) provide a trade space for customizing optimization to a particular problem or stimulation effect size. These simulations were directly based on empirical observations in humans, potentially increasing the likelihood of their replication *in vivo*.

UCB1’s relative superiority over other algorithms, particularly the more sophisticated parametric approach of Thompson sampling, may initially be surprising. This result is likely an effect of the blocked stimulation paradigm we emphasize, where stimulation parameters are held constant for some number of trials, and where the total number of samples is highly constrained. In addition to allowing the filtered variable *x_base_* to reach steady state, the blocked approach averages out stochastic noise, meaning that the RT or state estimate for each block is a reliable reporter of stimulation’s true effect. In the presence of reliable information/lowered noise, UCB1 and even greedy algorithms can outperform parametric approaches, because they do not require sufficient trials to reliably estimate a parametric distribution. In another scenario, with single trial measurements and more rapid changes in stimulation, TS might be superior.

This approach may address the target engagement problem that has limited the use of DBS in non-motor applications[2,19]. In the most straightforward example, the framework described above could be used to optimize VCVS DBS for depression or OCD. Current practice asks patients to rate their subjective mood as a clinician tests different contacts/settings[15,16]. In our proposed framework, patients would instead perform the MSIT or a similar cognitive task. Our prior work shows that patients can tolerate doing so for several hundred trials if given adequate breaks[36,37]. Even if the ensemble approach is needed, the current standard in VCVS DBS optimization is a 2+ day office procedure taking 8-10 hours [15]. Our proposed approach could actually take less time. Further, because task performance and task-based optimization could easily be performed in a clinical/office setting, it would require little change to the clinical workflow and no specialized equipment. Verification of target engagement through this task-based paradigm may lead to more robust clinical trial designs for new DBS indications. For instance, the algorithms above all can identify stimulation sites that do not improve RT/*x_base_*. Stimulation settings that do and do not improve cognitive control could be tested against each other in an ABAB or similar within-subjects design. Those designs have higher statistical power than the between-groups designs that failed in [54,55].

In the longer term, an objective assessment for behavioral target engagement could also inform surgical practice. For instance, multiple studies argue that clinical response to DBS, as measured with standard rating scales, is correlated with activation of specific white matter bundles[21,22,56,57]. This is likely also true of cognitive response – more dorsal fibers within the VCVS have larger behavioral effects[37], and specific cognitive domains are differentially linked to different sub-parts of the cortico-striato-thalamic circuitry that runs through VCVS[58]. By retrospective analysis, it may be possible to identify VCVS sub-bundles associated with the cognitive response described here, and to then target DBS electrodes specifically to those bundles.

The framework of optimizing DBS to engage a specific, objective, task-measured behavior may also lead to more robust animal models. The availability of rodent and non-human primate models for movement disorders DBS has enabled more rapid advances, because novel stimulation sequences and closed-loop paradigms can be more easily tested[59–61]. This would likely also be true for psychiatric DBS, but current animal models of mental disorders, grounded in human symptoms, have limited predictive power for clinical outcomes[62]. Cognitive assays, on the other hand, might be more reliably translated across species[63]. The method described here, using UCB1 to optimize between discrete stimulation sites, could equally be used to test among discrete types/paradigms of stimulation applied to the same electrode. Similar to [61], many different approaches might be compared sequentially in an animal cognitive assay, and the few reliable performers advanced to human pilots.

Conversely, the concept of combining a Bayesian optimizer with a filtering/ latent state model could be applied on longer timescales consistent with clinical outcomes, in both psychiatric disorders and neighboring applications such as chronic pain. Multiple authors have argued that DBS is difficult to develop for non-motor applications because the clinical outcomes are self reports, which are overly influenced by expectancy, environmental events, and other non-structured variations [2,19,20,64–66]. If self report could be made more reliable, there are case reports of successful neurostimulator programming through optimization algorithms [42,67]. Non-DBS-related changes in self-report outcomes can be considered as stochastic noise, in principle similar to the variations in RT that we smooth out through the filtering described in equations (1) and (2). The same approach could be applied to patient self report, e.g. collecting multiple noisy ratings at dense timescales and filtering to identify the underlying “true” rating. In cases where there are known environmental influences (e.g., a patient whose pain is influenced by time of day or by emotional stressors), these could be modeled as inputs, similar to how we modeled interference effects through *x_conflict_*.

## Limitations

The largest limitation of the current system is that it can optimize only a single discrete parameter (stimulation site). In practice, clinically effective DBS requires finding the correct combination of site(s), frequency, pulse width, amplitude, and temporal patterning (e.g., cycling). Algorithms exist for optimizing these continuous-valued parameters [41], but efficient approaches to multivariate, discrete-continuous hybrid optimization are not yet developed. As more variables are simultaneously considered, the response space explodes combinatorially and it becomes impossible to converge in clinically feasible times. Stimulation parameters may need to be optimized sequentially by univariate algorithms, or multivariate optimization might be constrained, e.g. by stronger prior probabilities derived from prior patients.

The current approach does not consider adverse effects. DBS at VCVS is well documented to cause negative side effects, including immediate high anxiety[14,15,68] and slower-onset impulsivity and mania [13,69]. Optimization algorithms should avoid settings known to cause these adverse effects, but the optimizers described here have no way to represent extreme negative outcomes. They also do not provide feedback as to whether the difference between two sites can be considered statistically significant, which may be clinically important if more than one site is close to the global optimum. All of these factors would need to be re-considered in optimizing DBS for a different cognitive effect, presumably with a different behavioral assay. The underlying COMPASS toolkit is flexible and can model most standard tasks[43], but model structure would still need to be customized for each new application, as would the noise covariance assumptions. This poses a challenge for DBS targets beyond VCVS[22,70]. It is not yet known which tasks/assays can measure engagement at other targets, which limits the use of this approach as a general-purpose tool in psychiatric DBS.

More broadly, the current approach is an indirect read-out of DBS target engagement. DBS ultimately exerts its effects by changing neurophysiology [2,71]. Other approaches to DBS optimization in psychiatry focus on identifying stimulation parameters that directly produce a desired physiological change[34], or on responding to changes in a target biomarker to leverage the state dependence of DBS effects [18,26]. We cannot inform responsive DBS with our proposed approach, nor can we match the precision of neurophysiology. Those more advanced approaches, however, require correspondingly advanced inpatient monitoring and analytic capabilities. The method proposed here requires only commodity computer hardware, and thus may be more cost-effective and scalable.

## Conclusion

In summary, we have demonstrated a proof of concept system for psychiatric DBS programming and online optimization, using cognitive control as the measure of effective target engagement. This may lead to more reliable clinical response and/or more robust design for clinical trials of the specific VCVS target studied here. Future work will include extending optimization to multiple DBS parameters and handling of stimulation-induced adverse events. Above all, any potential clinical use would require optimization software that was tested for usability and design robustness[72], a critical next step for validating our proposed approach.

## Acknowledgements

This work was supported by the Minnesota’s Discovery, Research, and Innovation Economy (MnDRIVE) initiative, the Minnesota Medical Discovery Team on Addictions, and the National Institutes of Health (R01MH124687, R01NS120851). The opinions herein are the position of the authors, not any governmental body or funding agency.

## Disclosures

ASW, AY, and TIN have unlicensed intellectual property in the area of brain stimulation optimization, including patent applications related to the subject of this paper. TIN holds equity in StimSherpa, a company developing optimization methods for neuromodulation. All other authors declare no financial conflicts of interest.

## Code Availability

At the time of publication, all code and data required for performing the above-described simulations will be made available in a GitHub repository https://github.com/tne-lab/DBSParameterOptimization2022.git.

## Notes

https://github.com/tne-lab/DBSParameterOptimization2022.git

### Competing Interest Statement

Alik S Widge, Ali Yousefi, and Theoden I Netoff have unlicensed intellectual property in the area of brain stimulation optimization, including patent applications related to the subject of this paper. Theoden I Netoff holds equity in StimSherpa, a company developing optimization methods for neuromodulation. All other authors declare no financial conflicts of interest.

